# Dopamine D_2_ receptor agonists abrogate neuroendocrine tumour angiogenesis to inhibit chemotherapy-refractory small cell lung cancer progression

**DOI:** 10.1101/2024.11.07.622543

**Authors:** Sk. Kayum Alam, Anuradha Pandit, Li Wang, Britteny A. Thiele, Parvathy Manoj, Marie Christine Aubry, Charles M. Rudin, Ying-Chun Lo, Luke H. Hoeppner

**Author notes:** Corresponding Author: Luke H. Hoeppner, PhD The Hormel Institute, University of Minnesota 801 16^th^ Avenue NE Austin, MN 55912.

## Abstract

Small cell lung cancer (SCLC) is difficult to treat due to its aggressiveness, early metastasis, and rapid development of resistance to chemotherapeutic agents. Here, we show that treatment with a dopamine D_2_ receptor (D_2_R) agonist reduces tumour angiogenesis in multiple *in vivo* xenograft models of human SCLC, thereby reducing SCLC progression. An FDA-approved D_2_R agonist, cabergoline, also sensitized chemoresistant SCLC tumours to cisplatin and etoposide in patient-derived xenograft models of acquired chemoresistance in mice. *Ex vivo*, D_2_R agonist treatment decreased tumour angiogenesis through increased apoptosis of tumour-associated endothelial cells, creating a less favourable tumour microenvironment that limited cancer cell proliferation. In paired SCLC patient-derived specimens, D_2_R was expressed by tumour-associated endothelial cells obtained before treatment, but D_2_R was downregulated in SCLC tumours that had acquired chemoresistance. D_2_R agonist treatment of chemotherapy-resistant specimens restored expression of D_2_R. Activation of dopamine signalling is thus a new strategy for inhibiting angiogenesis in SCLC and potentially for combatting chemotherapy-refractory SCLC progression.

---

Lung cancer is the leading cause of cancer deaths worldwide^1^. Small-cell lung cancer (SCLC) accounts for 15-17% of lung cancer cases and ∼200,000 deaths each year globally^2,3^. Most newly diagnosed SCLC patients present with extensive-stage disease and initially respond to first-line chemotherapy but frequently develop drug resistance, resulting in a dismal five-year survival rate of only 7%^4,5^. SCLC has recently been categorized into three subtypes with unique transcriptional characteristics and therapeutic vulnerabilities based upon differential expression of the transcription factors *ASCL1* (SCLC-A; 51%), *NEUROD1* (SCLC-N; 23%), and *POU2F3* (SCLC-P; 7%)^6,7^, and a fourth subtype exhibits low expression of all three transcription factors with high inflammation and susceptibility to immune checkpoint inhibitors (SCLC-I; 17%)^6^. Molecularly targeted therapies have improved survival time of patients with non–small-cell lung cancer (NSCLC)^5^, but similar targeted treatments in SCLC have largely failed, and chemotherapy has remained standard of care for over three decades. Outcomes from chemotherapy are poor, with a median overall survival rate of 9.4 months^8,9^. Recent progress has been made improving overall survival of SCLC patients with US Food and Drug Administration (FDA)-approved immunotherapies, including the PD-L1 inhibitors atezolizumab and durvalumab^8,10,11^. For example, the addition of atezolizumab to platinum-based frontline chemotherapy extends median overall survival by 2 months^11^. Unfortunately, however, only a small subset of patients with extensive-stage SCLC experience deep and durable responses to immune checkpoint blockade, and reliable prognostic biomarkers to identify such potential responders do not currently exist. In spite of robust initial responses to first-line platinum-etoposide with or without immunotherapy, nearly all SCLC patients eventually relapse^8^. Consequently, developing improved treatment approaches for extensive-stage SCLC is an urgent unmet need.

The tumour microenvironment has recently emerged as an active promoter of cancer progression. A dynamic and reciprocal relationship between cellular components of the tumour microenvironment and cancer cells is established early in tumorigenesis. For example, tumour-associated endothelial cells respond to cues within the tumour microenvironment to promote tumour angiogenesis; the newly formed vessels supply the tumour with oxygenated blood and provide a provisional matrix capable of supporting additional vascular sprouting and tumour growth. Vascular endothelial growth factor (VEGF) A is an essential regulator of tumour angiogenesis. VEGF was initially discovered as “vascular permeability factor”, a tumour-secreted protein that potently promotes microvascular permeability^12^. Only later was it discovered separately as an endothelial mitogen^13^ essential for the development of blood vessels^14–16^. VEGF signals predominantly through VEGF receptor 2 (VEGFR2) to regulate endothelial cell function by activating downstream signalling, including phospholipase Cγ–mediated activation of the mitogen-activated protein kinase (MAPK)/extracellular-signal-regulated kinase-1/2 (ERK1/2) pathway and phosphatidylinositol 3′ kinase (PI3K)-induced stimulation of the AKT (protein kinase B) signalling cascade^17,18^. VEGF plays a crucial role in the development and homeostasis of the lung vasculature, and the lungs exhibit the highest level of VEGF gene expression among physiologically normal tissues^19^. In pathological conditions, the expression of VEGF and its receptors is frequently affected by the specific characteristics of the disease and its stage in progression^20^. VEGF levels are higher in SCLC patients than in healthy individuals^21,22^; correspondingly, increased serum VEGF levels are the only independent prognostic factor other than tumour stage in untreated SCLC patients^23^, as confirmed by Zhan and colleagues through a large review and meta-analysis of VEGF expression in lung cancer^24^. Increased secretion of VEGF by tumour cells and upregulation of VEGFR2 in SCLC promote tumour angiogenesis, which provides tumours with the blood supply necessary to grow and metastasize.

Inhibition of angiogenesis has improved progression-free survival in several human cancers, including NSCLC^25,26^. Bevacizumab, a humanized monoclonal antibody that binds all forms of VEGF-A, was FDA-approved for the treatment of NSCLC in 2006^27^ but has limited efficacy against SCLC when combined with chemotherapy^28–35^. Many other angiogenesis inhibitors, including thalidomide^36,37^ and a variety of small molecule tyrosine kinase inhibitors of VEGF receptors, such as sunitinib^38–41^, vandetanib^42^, cediranib^43^, sorafenib^44^, and nintedanib^45^, have failed, achieved modest responses, or caused an unacceptable degree of toxicity in SCLC. Anlotinib, an orally administered tyrosine kinase inhibitor of VEGFR and other growth factor receptors, was recently approved by the China FDA as a third-line therapy for Chinese patients with SCLC following clinical trials (ALTER 1202 study)^46,47^; however, to date no anti-angiogenic agents have received regulatory approval in the US for treatment of SCLC. Importantly, because SCLC lacks predictive biomarkers for response to VEGF inhibition, whether a subset of SCLC patients do respond to anti-VEGF treatment is unknown. Here, we seek to overcome this issue by manipulating the dopamine signalling pathway to inhibit angiogenesis, progression, and drug resistance in SCLC.

Dopamine is an important neurotransmitter in the central nervous system that is produced by sympathetic nerves that end on blood vessels. Dopamine acts on its target cells in a cAMP-dependent manner through its receptors, which belong to the G protein–coupled receptor superfamily and are broadly classified as D_1_ and D_2_ types^48^. The D_1_ class includes D_1_R and D_5_R, which increase intracellular cAMP upon activation^49^. The D_2_ types, including D_2_R, D_3_R, and D_4_R, inhibit intracellular cAMP^49^. D_2_R is expressed by a variety of cell types, including neurons, immune cells, endothelial cells, and endothelial progenitor cells^48^ and colocalizes with VEGFR-2; D_2_R/VEGFR-2 crosstalk mediates dephosphorylation of VEGFR2^50,51^. Dopamine and other D_2_R agonists bind to D_2_R expressed on the surface of endothelial cells to inhibit VEGF-mediated angiogenesis; they also completely block accumulation of tumour ascites and tumour growth in mice^52^. Conversely, increased angiogenesis, tumour growth, and VEGFR2 phosphorylation are observed in D_2_R knockout mice^53^. D_2_R agonists increase the efficacy of anti-cancer drugs in preclinical models of breast and colon cancer through their anti-angiogenic effect on tumour-associated endothelial cells^54^. Dopamine signalling effector molecules regulate lung cancer progression^55–58^. In a lung injury model, we showed that dopamine inhibits pulmonary oedema through the endothelial VEGF-VEGFR2 axis^59^. We also demonstrated that an FDA-approved D_2_R agonist, cabergoline, reduced NSCLC growth in mice and reported that a subset of NSCLC patients have increased endothelial D_2_R expression, suggesting potential precision oncology treatment strategies are possible^60^. Work by several laboratories clearly suggests that this strategy can be translatable for therapy in different diseases, including cancer^53,61,62^. Based on these collective prior observations, we sought to understand how dopamine signalling regulates therapy-refractory SCLC progression and to determine whether D_2_R agonist treatment can inhibit growth of chemotherapy-resistant SCLC by reducing angiogenesis.

## Results

### D_2_R agonists inhibit SCLC growth in a human xenograft model

To begin testing the hypothesis that D_2_R agonists inhibit SCLC growth by reducing tumour angiogenesis, we orthotopically injected luciferase-labelled human DMS-53 SCLC cells into the left thorax of SCID mice (**Fig. 1a**). Non-invasive luciferase imaging demonstrated that the SCLC tumours had established themselves within the lungs of mice by 7 days after injection of the DMS-53 cells. Starting on day 8, we injected vehicle or the D_2_R agonist quinpirole (10 mg/kg) intraperitoneally every other day for 13 days (i.e., until day 20). Luciferase imaging on day 21 showed that DMS-53 lung tumour growth decreased in mice treated with D_2_R agonist relative to that in vehicle-treated control mice (**Fig. 1b-c; Supplementary** Fig. 1), supporting the hypothesis that D_2_R agonists inhibit SCLC growth in an orthotopic mouse model of human SCLC.

**Figure 1:**
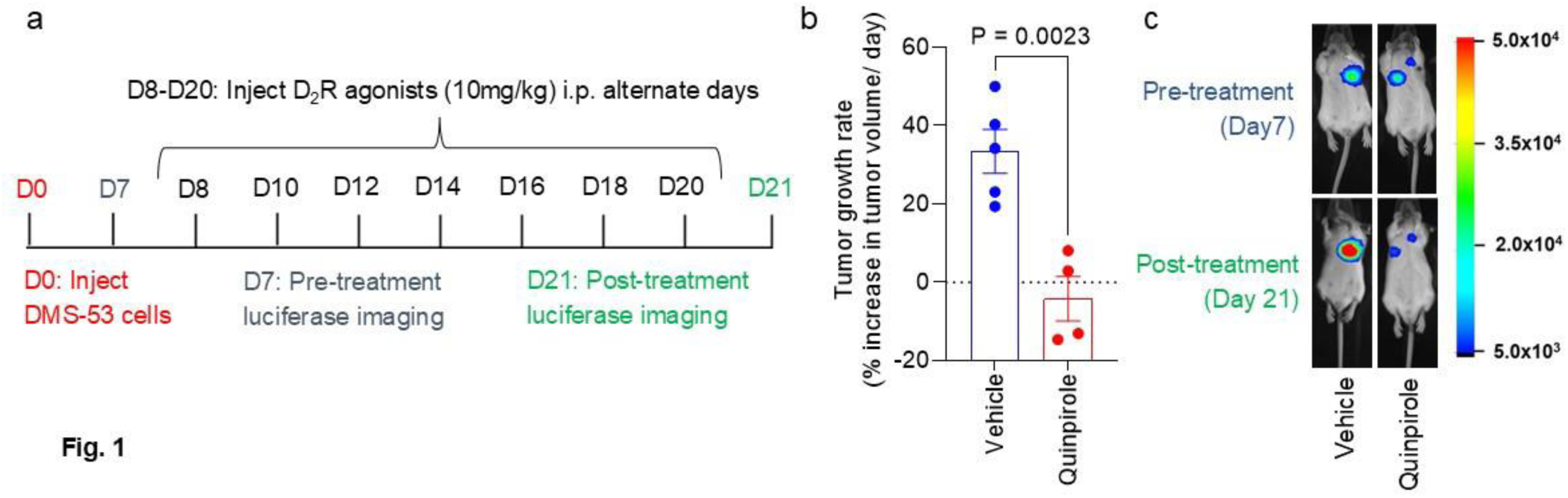
Activation of D_2_R signalling by quinpirole reduces tumour growth in a human small-cell lung tumour xenograft model. **a.** Experimental timeline: SCID mice were orthotopically injected with luciferase-labelled human DMS-53 SCLC cells, then imaged for bioluminescence 7 days after the injection but before the start of treatment. Mice then received intraperitoneal injections of PBS vehicle (control group) or 10 mg/kg quinpirole (treatment group) every other day for 13 days, after which they were imaged again for bioluminescence. **b.** Distribution of tumour growth rate across the vehicle- and quinpirole-treated groups. Each circle represents an individual mouse. Data are shown as mean ± SEM. A value of *P* ≤ 0.05 (two-way unpaired t-test) was considered significant. **c.** Representative luciferase imaging from day 7 (before treatment with D_2_R agonist) and day 21 (after treatment with D_2_R agonist).

### D_2_R agonist cabergoline reduces tumour growth in PDX models of SCLC

We next evaluated whether D_2_R agonists reduce neuroendocrine tumour progression using a human SCLC patient-derived xenograft (PDX) model. Specifically, a previously described human SCLC PDX specimen, named MSK-LX40^63^, was subcutaneously injected into nonobese severe combined immunodeficient γ (NSG) mice. After establishment of tumours of at least 100-200 mm^3^ in size, mice were intraperitoneally administered either vehicle (PBS) or cabergoline (5 mg/kg) daily for 2 weeks (**Fig. 2a**). Tumour growth was inhibited in SCLC PDX–bearing mice treated with cabergoline compared with growth in controls (**Fig. 2a**). At the endpoint, the mice were euthanized and their tumours harvested. Tumour weight and volume were lower in the cabergoline-treated group than in the controls (**Fig. 2b-d**). To test our hypothesis that D_2_R agonist treatment reduces SCLC tumour growth by reducing tumour angiogenesis and creating a less favourable tumour microenvironment that limits cancer cell proliferation, we co-stained tumour specimens harvested from the mice for CD31 to detect tumour-associated endothelial cells and for TUNEL (terminal deoxynucleotidyl transferase–mediated deoxyuridine triphosphate nick end labelling) to quantify apoptosis. As expected, we observed more CD31 and TUNEL co-staining in cabergoline-treated tumours than in vehicle-treated tumours (**Fig. 2e-f**), suggesting that D_2_R agonist treatment increases apoptosis of tumour-associated endothelial cells. *Ex vivo* immunostaining for Ki-67 revealed significantly less proliferation of cancer cells within the patient-derived SCLC xenografts of mice treated with cabergoline than of control mice treated with vehicle (**Fig. 2g-h**). Taken together, these findings suggest that the FDA-approved D_2_R agonist cabergoline decreases SCLC progression by reducing tumour angiogenesis through apoptosis of tumour-associated endothelial cells, leading to decreased proliferation of cancer cells within the SCLC PDX.

**Figure 2:**
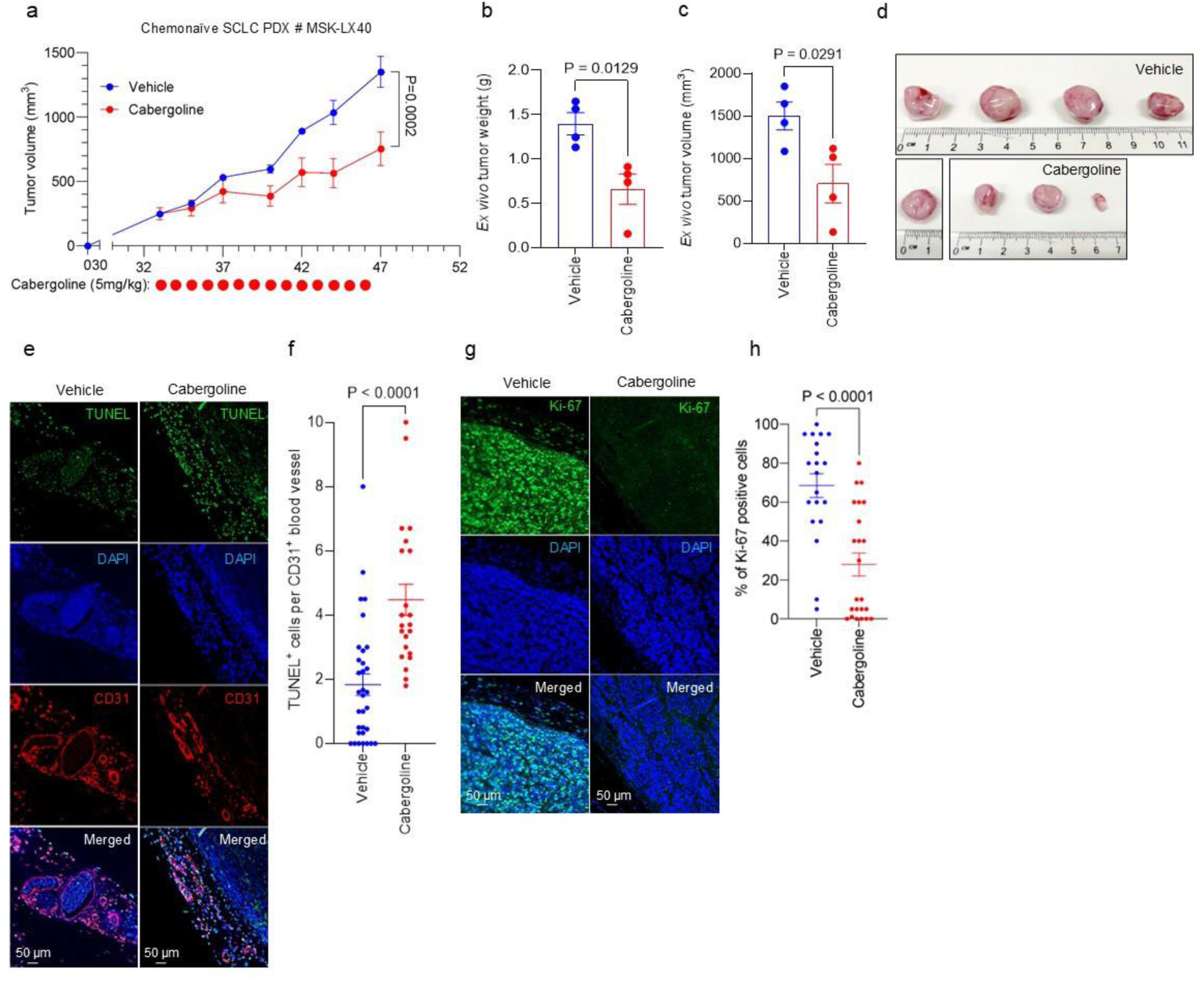
D_2_R agonist treatment reduces tumour growth and promotes apoptosis of tumour-associated endothelial cells in a chemonaïve human SCLC PDX model. **a-d**. NSG mice were subcutaneously injected with 5 × 10^6^ human SCLC cells originally derived from a chemonaïve patient (MSK-LX40)^63^. Mice were treated with either vehicle (10% DMSO in PBS) or cabergoline (5 mg/kg) daily for 14 days once mean tumour volume reached 100-200 mm^3^. Tumour growth was recorded by measuring tumour volume with callipers three times a week (a). Experiments were concluded before average tumour volume exceeded 1500 mm^3^. At the endpoint of the experiments, mice were euthanized, and xenografted tumours were harvested and weighed using a digital balance (b). Tumour volume was calculated from calliper measurements after extirpation (c). Each circle represents an individual mouse (b-c). Images of harvested tumours were acquired (d). **e**. Co-immunofluorescence staining was performed on formalin-fixed, paraffin-embedded (FFPE) tissues harvested from NSG mice (n=3 mice per group) treated with either vehicle or cabergoline to detect both the TUNEL^+^ and CD31^+^ cells in the tumour microenvironment. **f**. The colocalization of TUNEL and CD31 staining was quantified by counting the number of double-positive cells (i.e., TUNEL^+^ and CD31^+^ cells) per CD31^+^ blood vessel. Each circle represents one visual field (≥5 visual fields were counted for each tissue). **g**. Immunofluorescence staining was performed using a monoclonal Ki-67 antibody on FFPE tumour tissue specimens harvested from chemonaïve human SCLC tumour–bearing mice treated with vehicle (left) or 5 mg/kg cabergoline (right). **h**. The proliferation of tumour tissues harvested from cabergoline- and vehicle-treated mice (n=3 for each group) was quantified by counting the number of Ki-67-positive cells per visual field (≥5 visual fields were counted for each tissue). Data are shown as mean ± SEM. A value of *P* ≤ 0.05 (two-way unpaired t-test) was considered significant.

### D_2_R agonist cabergoline shows anti-proliferative effects in PDX models of chemotherapy-refractory SCLC

Given that normalization of the tumour vasculature improves intratumoral accessibility of anti-cancer agents^64^, we hypothesized that D_2_R agonist treatment may reduce chemotherapy-refractory SCLC progression by decreasing tumour angiogenesis within the lung tumour microenvironment, enhancing the anti-cancer effects of the chemotherapy. To test this hypothesis, we relied on three SCLC PDXs (MSK-LX40R, JHU-LX108R, JHU-LX33R) that had been previously passaged through mice treated with the standard-of-care chemotherapy regimen, cisplatin + etoposide, such that the PDXs acquired chemoresistance^63^. We sought to evaluate the impact of D_2_R agonist treatment on mice bearing chemotherapy-refractory SCLC PDX tumours. Therefore, we implanted chemoresistant SCLC PDX tumours into the right flank of NSG mice, monitored animals every week until the establishment of the palpable tumour (i.e., tumour volume 100-200 mm^3^), and then randomized the mice into two groups. Each group of mice received weekly regimens of 5 mg/kg cisplatin on day 1 and 8 mg/kg etoposide on days 1, 2, and 3. One of the two groups also received weekly regimens of 5 mg/kg cabergoline on days 1-5 of each week for 2 to 5 weeks (depending on how aggressive each PDX was). The combination treatment of cabergoline and chemotherapy reduced tumour growth across all three PDX models of chemorefractory SCLC (**Fig. 3a-c**). At the conclusion of treatment, the mice were euthanized and tumours excised, weighed, and measured. As expected, we observed a decrease in *ex vivo* tumour weight and volume in cabergoline-treated mice compared with those treated without cabergoline (**Fig. 3d-l**), suggesting that D_2_R agonism sensitizes chemotherapy-refractory human SCLC to cisplatin and etoposide.

**Figure 3:**
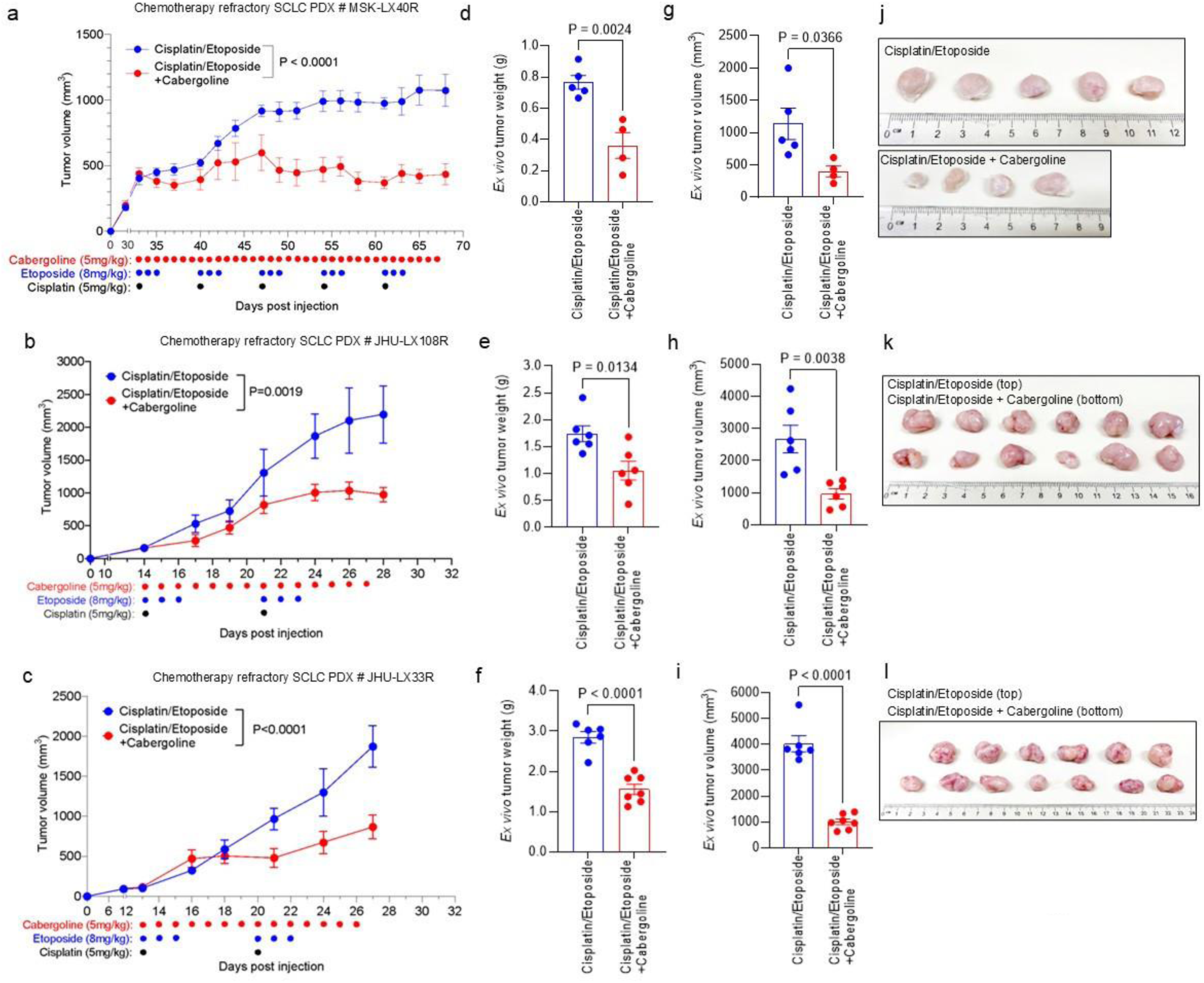
Treatment with D_2_R agonist and chemotherapy overcomes chemoresistance in human SCLC PDX models. **a-c.** Five million human tumour cells harvested from chemoresistant SCLC PDXs (MSK-LX40R [a], JHU-LX108R [b], and JHU-LX33R [c]) were subcutaneously injected into the right flanks of NSG mice. When tumour volume reached 200 mm^3^ in size, mice were randomly divided into treatment groups such that each group had similar mean tumour volumes. Mice were treated by intraperitoneal injection of 5 mg/kg cisplatin on day 1 and 8 mg/kg etoposide on days 1, 2, and 3 with or without 5 mg/kg cabergoline daily for the indicated times. Tumour volume was measured 2-3 times per week until the final tumour reached either five times the initial tumour volume (a) or 2000 mm^3^ in size (b-c). **d-f.** At the endpoint, the extirpated tumours were weighed on a digital scale. **g-i.** The final volume of the tumours from euthanized mice was measured using a digital calliper. **j-l.** Harvested tumours from each group were arranged randomly, and pictures were taken of the tumours and a clear plastic ruler with a phone camera. Each circle represents a tumour harvested from an individual mouse. Data are shown as mean ± SEM. A value of *P* ≤ 0.05 obtained with two-way ANOVA followed by Sidak’s multiple test (a-c) or two-way unpaired t-test (d-i) was considered significant.

To evaluate whether D_2_R agonist treatment promotes apoptosis of tumour-associated endothelial cells, we performed *ex vivo* co-immunofluorescence staining with TUNEL and anti-CD31 antibodies using SCLC tissue specimens derived from mice harbouring the chemoresistant MSK-LX40R PDX. More endothelial cells were TUNEL-positive in cabergoline- and cisplatin/etoposide (C/E)-treated tissue specimens than in samples treated only with C/E (**Fig. 4a-b**), suggesting that activation of D_2_R signalling stimulates apoptosis in tumour-associated endothelial cells. Given that normalization of the tumour vasculature improves intratumoral accessibility of anti-cancer therapies^64^, we hypothesized that D_2_R agonist treatment may reduce chemotherapy-refractory SCLC progression. Indeed, histological analysis of the MSK-LX40R model revealed that mice treated with cabergoline and C/E had lower Ki-67 expression than mice treated exclusively with C/E (**Fig. 4c-d**).

**Figure 4:**
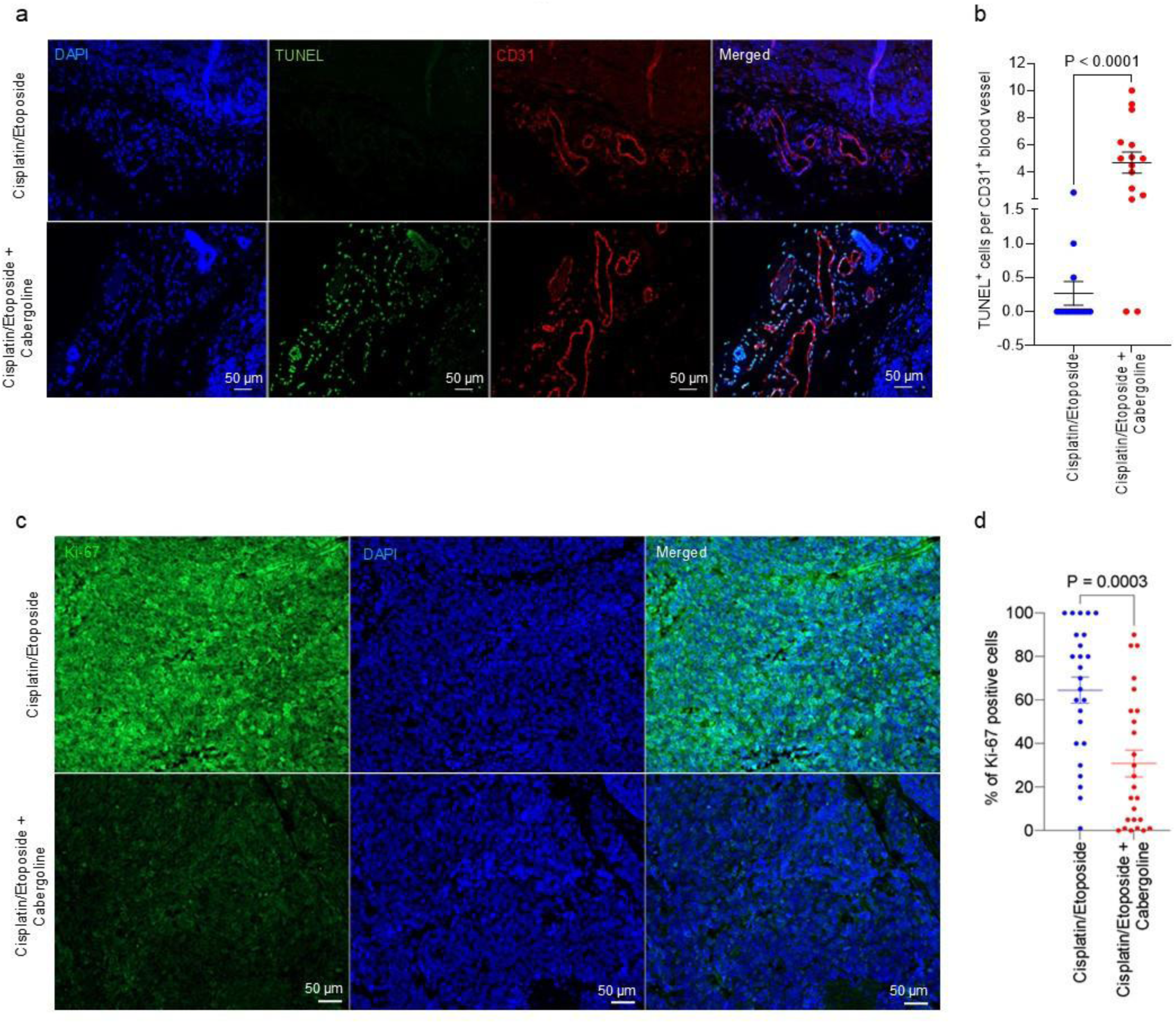
D_2_R agonist treatment together with chemotherapy decreases tumour cell proliferation and promotes endothelial apoptosis in a chemoresistant SCLC PDX model. **a.** FFPE tumour tissues (n=3 per group) treated with either chemotherapy alone or both cabergoline and chemotherapy were used for a co-immunofluorescence study to detect double-positive TUNEL^+^ and CD31^+^ cells in the tumour microenvironment. **b.** To measure D_2_R agonist-induced apoptosis in tumour-associated blood vessels, colocalization of TUNEL and CD31 staining was quantified in FFPE tumour tissues (n=3 per group) harvested from mice bearing MSK-LX40R SCLC PDXs. Each circle represents one visual field, and five visual fields were counted for each tissue. **c.** Immunofluorescence using monoclonal Ki-67 antibody was performed on FFPE tumour tissues (n=5 per group) obtained from a chemoresistant human SCLC PDX model (MSK-LX40R) treated with 5 mg/kg cisplatin on day 1 and 8 mg/kg etoposide on days 1, 2, and 3 with or without 5 mg/kg cabergoline daily. Scale bar, 50 µm. **d.** The number of Ki-67-positive cells from the experiment illustrated in (c) was quantified. Each circle represents one visual field, and five visual fields were counted for each tissue. Data are shown as mean ± SEM. A value of *P* ≤0.05 was considered significant, two-way unpaired t-test.

To further test our hypothesis, we first created 3D SCLC PDX organoids from JHU-LX33R (**Fig. 5a**). We next stably transduced human umbilical vein endothelial cells (HUVECs) with either a D_2_R-specific shRNA to silence D_2_R or a control LacZ-specific shRNA and treated these HUVECs with a D_2_R agonist (**Supplementary** Fig. 2). We collected the conditioned medium from the D_2_R agonist–treated HUVEC cultures and placed it on the dissociated SCLC PDX organoids for 72 hours (**Fig. 5a**). Conditioned medium derived from D_2_R agonist– treated HUVECs stably transduced with the control LacZ shRNA elicited a higher apoptotic response of PDXs than vehicle-treated shControl HUVEC (**Fig. 5b; Supplementary** Fig. 3). As expected, conditioned medium derived from D_2_R agonist–treated HUVECs stably transduced with D_2_R shRNA did not cause a change in apoptosis relative to vehicle-treated shControl HUVEC (**Fig. 5b; Supplementary** Fig. 3), suggesting that D_2_R agonist signalling through D_2_R in endothelial cells is necessary to produce growth factors and cytokines present in the conditioned medium that stimulate apoptosis of the SCLC PDX cells. These observations correspond with our *in vivo* findings demonstrating that D_2_R agonist treatment reduces chemotherapy-refractory SCLC progression.

**Figure 5:**
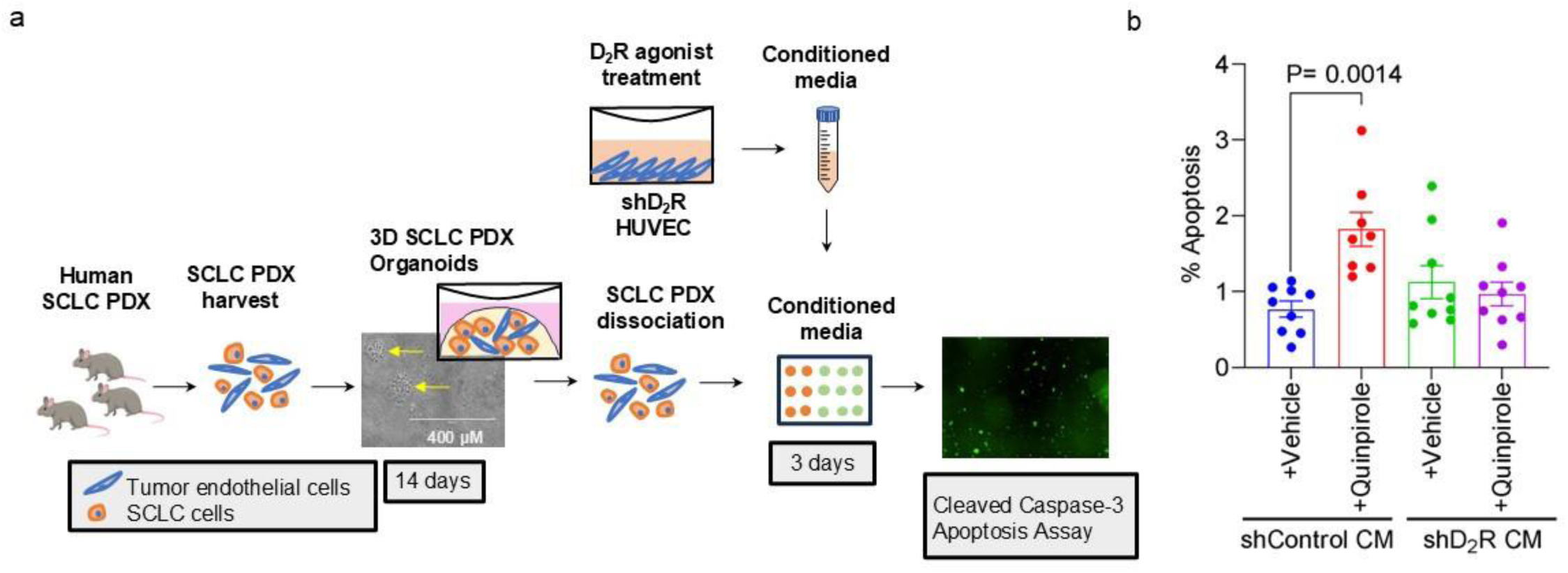
Conditioned medium from human endothelial cells treated with a D_2_R agonist promotes apoptosis of SCLC organoids. **a.** The SCLC PDX JHU-LX33R was suspended in Matrigel and plated at a density of 1 × 10^5^ cells in 50 µL. Complete growth media was added to the PDXs, which were grown in a humidified chamber at 37 °C supplied with 5% CO_2_. After 14 days, the Matrigel domes were dissolved by adding dispase (1 U/mL) containing cold complete medium, followed by TrypLE, then the cells were replated at a density of 10^4^ cells per well of a 96-well plate in Matrigel and topped off with 100 µL conditioned medium harvested from HUVECs treated with a D_2_R agonist (quinpirole). **b.** Conditioned media collected from HUVECs treated with the D_2_R agonist quinpirole (50 µM) increased the caspase-3–mediated apoptotic response in the SCLC chemotherapy-resistant organoid model.

### Chemotherapy-resistant SCLC-A specimens express less D_2_R on the surface of tumour-associated endothelial cells than matched chemotherapy-naïve specimens

We previously demonstrated a positive correlation between endothelial D_2_R expression and tumour stage through immunostaining of tumour specimens from NSCLC patients^60^. Therefore, we sought to assess D_2_R protein expression by immunostaining in paired chemonaïve and chemoresistant specimens from a cohort of SCLC-A patients (**Supplementary Table 1**). Briefly, tumour specimens were collected from SCLC patients before chemotherapy (i.e., chemonaïve) and following the development of chemotherapy-refractory disease progression. Following immunostaining with a D_2_R-specific antibody, a pulmonary pathologist (Y-C.L.) reviewed the D_2_R immunohistochemistry staining and scored the percentage of D_2_R-positive endothelial cells present in each stained lung tissue specimen. The number of D_2_R-positive endothelial cells in tumour specimens obtained following chemotherapy resistance was lower than that in the paired samples biopsied prior to chemotherapy treatment (**Fig. 6**). Collectively, our results suggest that protein expression of D_2_R decreases as SCLC-A tumours acquire resistance to chemotherapy.

**Figure 6:**
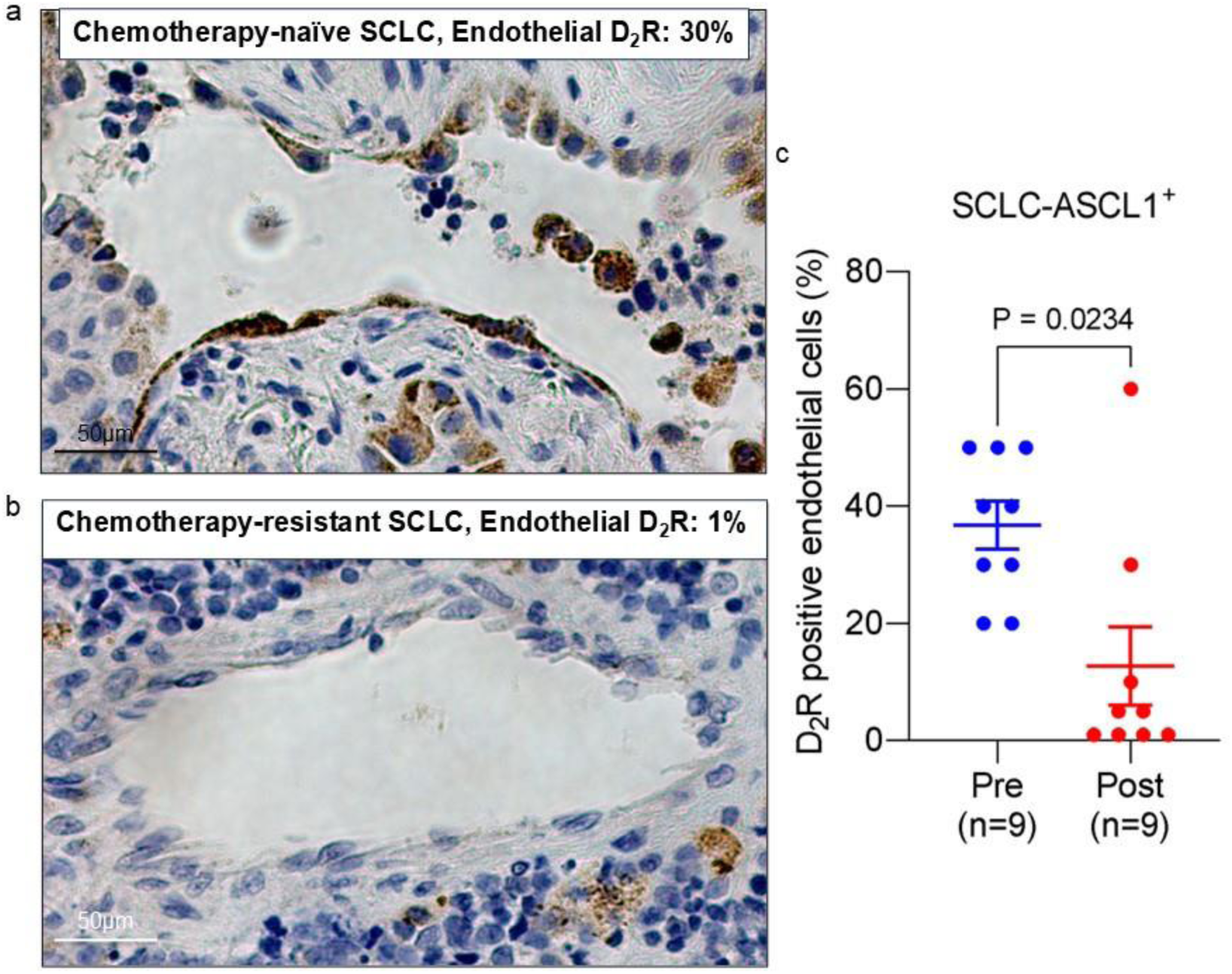
D_2_R is expressed by tumour-associated endothelial cells derived from SCLC patients. **a-b.** Immunohistochemistry was performed using a monoclonal D_2_R antibody on FFPE human SCLC-A tissue samples biopsied from each patient before the start of the chemotherapy (i.e., chemonaïve) and following progressive disease after chemotherapy (i.e., chemoresistant). Representative images from paired chemonaïve (a.) and chemoresistant (b.) SCLC tumour specimens used to assess D_2_R expression in tumour-associated endothelial cells before and after the chemotherapy. **c.** Each tissue was scored for the percentage of endothelial cells that were D_2_R-positive, and each circle on the graph represents an individual tissue (n=9). Data are shown as mean ± SEM. A value of *P* ≤ 0.05 (two-way unpaired t-test) was considered significant.

### Endothelial expression of D_2_R increases in chemoresistant SCLC PDX models in response to D_2_R agonist treatment

Because D_2_R expression on the surface of SCLC tumour-associated endothelial cells decreases as the tumours developed resistance to chemotherapy, we evaluated whether D_2_R agonist treatment affected the expression patterns of D_2_R in this context. Specifically, chemotherapy-sensitive MSK-LX40 or chemotherapy-resistant MSK-LX40R SCLC-A PDX specimens were subcutaneously implanted into the right flank of NSG mice. The mice were monitored until the establishment of palpable tumours (i.e., tumour volume 100-200 mm^3^). Mice were then randomized into four groups. Two groups received weekly regimens of 5 mg/kg cisplatin on day 1 and 8 mg/kg etoposide on days 1, 2, and 3. One of these two groups also received weekly regimens of 5 mg/kg cabergoline on days 1-5 of each week for three weeks. The third group received this cabergoline regimen in the absence of C/E. The fourth group of mice were administered vehicle as a negative control. The combination treatment of cabergoline and C/E reduced tumour growth relative to either treatment alone or vehicle control (**Fig. 7a-c**). An immunofluorescence-based colocalization study showed colocalization of D_2_R and CD31, indicating that endothelial cells express D_2_R in both MSK-LX40 and MSK-LX40R tumour tissues, although endothelial D_2_R expression was lower in vehicle-treated chemotherapy-resistant MSK-LX40R tumour tissues than in vehicle-treated chemotherapy-sensitive MSK-LX40 specimens (see vehicle-treated group in **Fig. 7d**; **Supplementary** Fig. 4), similar to our observation in paired chemonaïve and chemoresistant specimens from the cohort of SCLC-A patients (**Fig. 6**). Our immunofluorescence results showed no statistically significant changes in endothelial D_2_R expression upon C/E treatment between chemonaïve and chemoresistant SCLC PDXs (see C/E-treated group in **Fig. 7d**; **Supplementary** Fig. 4). However, cabergoline treatment alone or together with C/E was associated with a statistically significant increase in endothelial D_2_R expression in chemoresistant MSK-LX40R tumour tissues compared to chemonaïve tissue samples (see cabergoline- and C/E & cabergoline-treated group in **Fig. 7d**; **Supplementary** Fig. 4). Our collective results suggest that while lung endothelial D_2_R expression decreases as SCLC develops acquired resistance to chemotherapy, administration of the D_2_R agonist cabergoline results in increased expression of D_2_R, suggesting that low D_2_R expression in chemotherapy-refractory SCLC will not necessarily render D_2_R agonist treatment ineffective.

**Figure 7:**
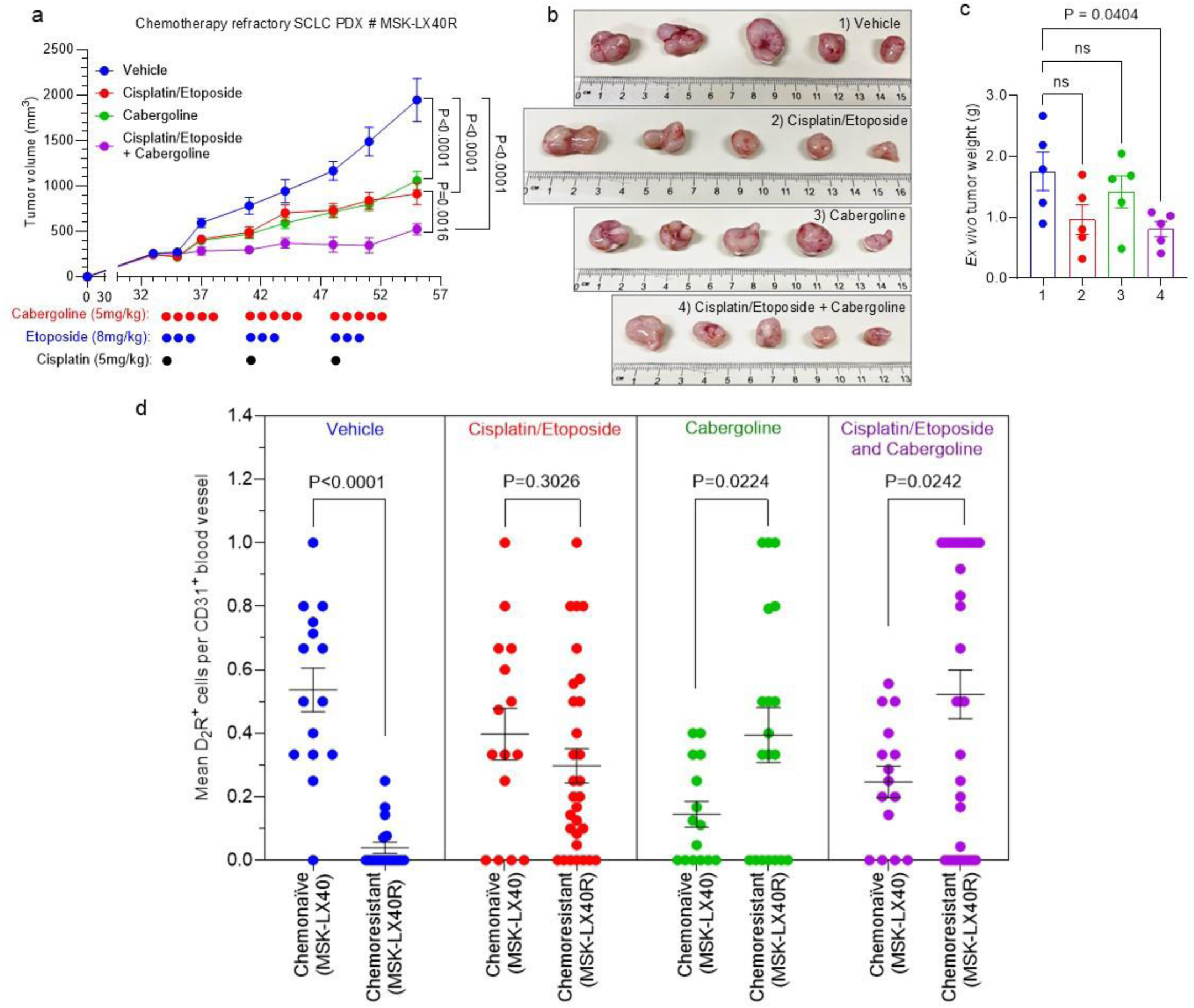
Endothelial D_2_R expression is increased by D_2_R agonist treatment in a chemoresistant SCLC PDX model. **a-c.** NSG mice were subcutaneously implanted with 5 × 10^6^ cells obtained from chemoresistant (MSK-LX40R) human SCLC PDXs. Mice were randomly divided into four groups: 1) vehicle; 2) cisplatin/etoposide; 3) cabergoline; and 4) cabergoline & cisplatin/etoposide. When mean tumour volume reached 200 mm^3^, mice were intraperitoneally administered vehicle (10% DMSO in PBS); 5 mg/kg cisplatin on day 1 and 8 mg/kg etoposide on days 1, 2, and 3, with or without 5 mg/kg cabergoline; or 5 mg/kg cabergoline alone five times a week. Tumour volume was measured three times a week until the vehicle-treated final tumour volume reached 2000 mm^3^ in size (a). At the endpoint, tumours were harvested from euthanized mice and photographs were taken to visualize gross morphology (b). Weight of the extirpated tumours was recorded using a digital scale (c). **d.** Co-immunofluorescence staining was performed on FFPE tumour tissues (n=3 per group) using primary antibodies against CD31 and D_2_R. The number of endothelial cells expressing D_2_R was quantified by counting the number of double-positive cells (i.e., D_2_R^+^ and CD31^+^) divided by the CD31^+^ blood vessels present in a single visual field (≥5 visual fields were counted for each tissue). Data are shown as mean ± SEM. A value of *P* ≤ 0.05 (two-way unpaired t-test) was considered significant.

## Discussion

In this study, we sought to overcome the ineffectiveness of angiogenesis inhibitors for SCLC treatment by manipulating the dopamine signalling pathway to inhibit angiogenesis, progression, and drug resistance in human SCLC. We demonstrated that D_2_R agonists sensitize chemoresistant SCLC tumours to C/E using several PDXs to model acquired chemoresistance in mice. Biochemical analyses of tissue from these mice suggest that treatment with a D_2_R agonist reduces tumour angiogenesis through increased apoptosis of tumour-associated endothelial cells, leading to a less favourable microenvironment for tumours and impeding cancer cell proliferation. In paired specimens derived from individuals with SCLC-A subtype tumours before and after the progression of chemotherapy-refractory disease, we showed that tumour-associated endothelial cells express D_2_R before exposure to chemotherapy. As SCLC tumours develop chemoresistance, D_2_R levels decrease, but treatment with a D_2_R agonist enhances D_2_R expression in endothelial cells from chemotherapy-resistant specimens.

Tumour angiogenesis is induced by VEGF-A binding to VEGFR2 on the surface of tumour-associated endothelial cells to activate downstream signalling pathways. Therefore, VEGF-A is a therapeutic target for inhibition of angiogenesis and normalization of tumour vasculature^65,66^. Bevacizumab, a humanized anti-VEGF-A monoclonal antibody, was approved by the FDA for the treatment of NSCLC in 2006^27^ and for the treatment of recurrent glioblastoma in 2009^67^. Moreover, bevacizumab treatment has been linked to inhibition of vessel growth, regression of newly formed vessels, and normalization of the vasculature to facilitate the delivery of cytotoxic chemotherapy^65^. Bevacizumab in combination with C/E chemotherapy has been shown to improve progression-free survival in a selected SCLC patient population, but no significant improvement of overall survival benefits was observed^68^. Similarly, the combination of bevacizumab and paclitaxel in chemosensitive SCLC has failed to yield any noteworthy clinical outcomes^69^, underlining the fundamental differences between SCLC and NSCLC^70^. The failure of anti-angiogenic agents to improve overall survival in SCLC underscores the need for finding an effective targeted therapy to improve the outcome of chemotherapy and/or anti-angiogenic therapy in relapsed chemorefractory SCLC.

Our rationale for assessing the anti-cancer and anti-angiogenesis effects of D_2_R agonists as a potential therapy to inhibit human SCLC progression stems from (1) our prior finding that D_2_R agonists inhibit angiogenesis and reduce tumour progression in murine models of NSCLC^60^, (2) the role of dopamine signalling in physiological neuronal function^71^, and (3) the neuroendocrine origin of SCLC^72^. In support of our hypothesis that D_2_R agonists inhibit SCLC growth by reducing tumour angiogenesis, a meta-analysis of 348,780 patients with Parkinson’s disease showed that patients who received dopaminergic therapy for the Parkinson’s had a 47% reduction in risk of developing lung cancer^73^. Furthermore, the FDA-approved D_2_R agonist cabergoline reduces tumour size in prolactinoma, a non-cancerous adenoma of the pituitary gland which typically has high D_2_R expression^74,75^.

Identifying biomarkers that predict responsiveness to therapy beyond classification of the tumour into one of the four current subtypes of SCLC will be critical to advancing treatment. Schlafen family member 11 (SLFN11) is a factor implicated in DNA damage repair deficiency. SCLC-A cell lines with high expression of SLFN11 are more resistant to cisplatin, whereas SLFN11 low expression was accompanied by increased sensitivity to cisplatin^63^, suggesting a potential clinical biomarker. Similarly, our observation that dynamic changes in D_2_R expression in tumour-associated endothelial cells depend on tumour responsiveness to chemotherapy and the activation status of D_2_R signalling may lead to a clinically useful biomarker.

Dopamine and D_2_R agonists have been shown to selectively inhibit VEGF-induced angiogenesis and vascular permeability by negatively regulating VEGFR2 phosphorylation^52^, resulting in the inhibition of endothelial cell migration^51^. Specifically, dopamine treatment increases VEGF-induced phosphorylation of phosphatase-2 containing Src homology region 2 domain (SHP-2) and its phosphatase activity in HUVECs. Subsequent dephosphorylation of VEGFR2 at Y951, Y996, and Y1059 by active SHP-2 inhibits VEGF-dependent signalling events, including those that promote angiogenesis and endothelial cell migration^51^. For example, in both dopamine-depleted and D_2_R-knockout mice, VEGF-induced phosphorylation of VEGFR2, MAPK, and focal adhesion kinase is substantially increased relative to the levels in control mice, indicating that dopamine signalling through D_2_R regulates these signalling pathways required for endothelial cell barrier integrity, proliferation, and migration^76^. Correspondingly, dopamine treatment in endothelial progenitor cells prevents their participation in tumour neovascularization by inhibiting their mobilization from the bone marrow niche^61^. Prior work provides a strong rationale for the concept that D_2_R agonist-mediated inhibition could be an effective therapeutic strategy^53,61,62^. Studies have demonstrated that disrupting peripheral dopaminergic nerves promotes tumour growth by triggering VEGF-dependent angiogenesis^53^, whereas dopamine treatment reduces the migration of tumour-promoting endothelial progenitor cells from the bone marrow^61^. D_2_R agonists have been shown to enhance the effectiveness of anti-cancer drugs in preclinical models of breast and colon cancer^54^.

Efforts to study the biology of SCLC have been hampered by the fact that most cases are inoperable, and biopsies are rarely obtained at recurrence. We sought to overcome this hurdle by relying on several previously generated PDX models of human SCLC resistance to C/E in mice^63^. Moreover, we have established an SCLC organoid model derived from the SCLC PDX tissue to study the effects of D_2_R agonists on functional processes, such as apoptosis, that could reduce chemotherapy-refractory SCLC progression. We have also taken advantage of paired SCLC-A subtype tumour specimens obtained from individual patients prior to chemotherapy (i.e., chemotherapy naïve/sensitive) and following chemotherapy-refractory disease progression (i.e., chemotherapy resistant). We anticipate that these models and specimens will drive future efforts to study how activation of D_2_R signalling promotes anti-angiogenic responses in the tumour microenvironment, particularly focusing on how cancer-associated fibroblasts, immune cells, stromal cells, and endothelial cells alter the tumour microenvironment to regulate tumour cell function.

Our studies primarily used immunocompromised mice to study the role of dopamine signalling in human SCLC progression, and the inability to model the contributions of the immune system is a limitation of our work. Future research using genetically engineered mouse models of human SCLC are needed to better understand how D_2_R agonist treatment affects immunoregulation within the tumour microenvironment.

Recent pursuits to better understand the molecular basis of chemotherapy-refractory SCLC progression have primarily focused on the contributions of tumour cell–intrinsic factors. Here, we highlight the importance of cancer cell–extrinsic regulation of the tumour microenvironment by demonstrating that activation of D_2_R signalling in tumour-associated endothelial cells by D_2_R agonists inhibits tumour angiogenesis and reduces chemotherapy-refractory SCLC growth. In accordance with our findings, dopamine, upon binding to D_2_R, reduces stress-mediated ovarian cancer growth by inhibiting tumour angiogenesis and stimulating tumour cell apoptosis^77^. Although it is well known that anti-angiogenesis therapies disrupt the tumour vasculature, the D_2_R agonist-mediated anti-angiogenesis process can also transiently “normalize” the abnormal structure and function of tumour vasculature to make it more efficient for oxygen and drug delivery^64^, which was clearly seen in our preclinical models (**Figs. 1-4**). Similar effects of dopamine on increased uptake of 5-fluorouracil have also been reported in human HT29 xenograft mouse models of colorectal cancer^78^. Our results are supported by the reported anti-tumour effects of endothelial D_2_R activation by dopamine in various solid tumours^49,52,60,76^. Although beyond the scope of these studies and not directly tested, we speculate that D_2_R agonists may increase transient normalization of tumour vessels, thereby producing a temporary increase in oxygen, alleviating hypoxia, and increasing the efficacy of conventional chemotherapies. Furthermore, we speculate that D_2_R agonist treatment may reduce immunosuppression within the tumour microenvironment, based upon our prior studies demonstrating that D_2_R agonist treatment reduces tumour-infiltrating myeloid-derived suppressor cells in NSCLC^60^. Furthermore, inhibition of VEGF signalling has been shown to stimulate CD4^+^ and CD8^+^ T cell activation and tumour infiltration, thereby reprogramming the tumour microenvironment^79^. In conjunction with immune checkpoint inhibitors, anti-angiogenic drugs can sensitize cancer cells to treatment. Other neuroendocrine tumours, RT2-PNET (pancreatic neuroendocrine tumour) and mammary carcinoma (MMTV-PyMT), showed successful treatment with a combination of anti-VEGFR-2 and anti-PD-L1 antibodies, inducing formation of high endothelial venules that facilitate enhanced infiltration of cytotoxic lymphocytes, activity, and tumour cell destruction^80^. While future research is necessary, D_2_R agonist treatment likely helps reprogram the immunosuppressive tumour microenvironment to enhance immune responses through reduction of tumour-infiltrating myeloid-derived suppressor cells and stimulation of T cells, making the tumour highly susceptible to enhanced anti-cancer responses through immune checkpoint inhibitors like anti-PD1/CTLA4.

## Methods

### Cell culture

The human SCLC cell line DMS-53 was purchased from The European Collection of Authenticated Cell Cultures (Salisbury, UK) via Sigma-Aldrich. This cell line was maintained in RPMI-1640 medium (Corning) supplemented with 10% foetal bovine serum (FBS; Millipore) and antibiotics. Human embryonic kidney (HEK) 293T cells purchased from American Type Culture Collection were cultured in Dulbecco’s modified Eagle’s medium (DMEM; Corning) supplemented with 10% FBS (Millipore),1% penicillin/streptomycin antibiotics (Corning), and 25 μg/mL plasmocin (Invivogen). Human umbilical vein endothelial cells (HUVECs; Lonza) were cultured in endothelial cell growth basal medium (EBM; CC-3121, Lonza) supplemented with the Endothelial Cell Growth Medium SingleQuots (CC-4143, Lonza). HUVECs of passages four to five were used and cultured in plates coated with bovine collagen type I protein (354231, Corning). All cell lines were grown in a humidified chamber at 37°C supplied with 5% CO_2_. Cell lines were authenticated by their source at the time of purchase and were subsequently routinely authenticated via morphologic inspection.

### Organoid culture

SCLC PDXs were suspended in growth factor–reduced Matrigel (Corning) and plated at a density of 10^5^ cells into 50 µL droplets in each well of a 24-well plate. Once the droplets solidified, complete growth medium consisting of DMEM supplemented with 10% FBS, 1% penicillin/streptomycin antibiotics (Corning), and 25 µg/mL Plasmocin (Invivogen) was added to the PDXs. Cells were grown in a humidified chamber at 37 °C supplied with 5% CO_2_ for 14 days. On the 14^th^ day, medium from each well was removed, and the Matrigel domes were dissolved by adding dispase (1 U/mL) containing cold complete media. Following incubation at 37°C for 90 mins with gentle agitation, the cells were centrifuged at 433 × g for 5 minutes. TrypLE (Gibco) was added to the pellet and incubated at 37°C for 10 mins. Following centrifugation at 433 × g for 5 minutes, the cells were replated at a density of 10^4^ cells per well of a 96-well plate in Matrigel and topped off with 100 µL media.

### Cleaved caspase-3 assay

Collagen at a concentration of 20 µg/mL was used to coat 10-cm plates. HUVECs were seeded at a density of 6 × 10^5^ cells and were grown to 70% confluency. Quinpirole and cabergoline were added at concentrations of 50 µM and 100 µM, respectively. After 72 hours, conditioned media were collected from the cells, filtered, and added to PDXs along with the Caspase-3 green dye (Sartorius) and BioTracker nuclear red dye (Sartorius). The plates were placed in IncuCyte for up to 96 hours with images taken every 24 hours.

### Generation of stable cell lines

Control pGIPZ-shScramble and lentiviral pGIPZ-shD_2_R plasmids designed to silence D_2_R expression were purchased from the University of Minnesota Genomics Center. Transient transfection of shRNA plasmids along with their corresponding packaging plasmids was performed in 293T cells using Effectene transfection reagent (Qiagen) in accordance with the manufacturer’s protocol. Lentivirus was collected from the cell culture medium at 48 and 72 h after transfection. Collected medium was then passed through a 0.45-µm syringe filter (Millipore) to remove cell debris. One-third volume of Lenti-X concentrator (Takara) was added to the cell culture medium and incubated overnight at 4°C. Precipitated lentivirus was collected from the cell culture medium after a brief centrifugation at 2000 × g for 1 h at 4°C. HUVECs were transduced twice with concentrated lentivirus diluted in fresh EBM containing 10 µg/mL Polybrene (Millipore). HUVECs were grown in EBM containing 2 µg/mL puromycin (Sigma) for 72 h to select positive clones that were resistant to puromycin and expressed GFP. The knockdown of D_2_R was confirmed by western blotting.

Human DMS-53 SCLC cells were transduced with retrovirus containing luciferase genes. Briefly, MSCV Luciferase PGK-hygromycin plasmid was obtained as a gift from Dr. Scott Lowe through Addgene (https://www.addgene.org/18782/). Retroviral luciferase plasmids and their corresponding packaging plasmids were transfected in HEK-293T cells using Effectene transfection reagent (Qiagen) in accordance with the manufacturer’s protocol. Retro-X concentrator (Takara) was used to concentrate retrovirus using cell culture media collected at 48 h and 72 h after transfection. Concentrated retrovirus diluted in fresh RPMI 1640 medium containing 10 µg/mL Polybrene (Millipore) was added to growing DMS-53 cells. Seventy-two hours after transduction, 5 µg/mL hygromycin (Sigma-Aldrich) was added to the cell culture medium, and DMS-53 cells were incubated for additional 72 h. Luciferase-labelled stable human DMS-53 cells were confirmed based on the expression of luciferase and sustained growth in the presence of hygromycin.

### Antibodies and drugs

Primary antibodies were used in western blotting experiments to detect expression of D_2_R (Abcam, cat no.: ab85367; 1:1000) and α-tubulin (Santa Cruz Biotechnology, cat no.: sc-5386; 1:500). Horseradish peroxidase (HRP)-conjugated anti-rabbit (Cat no.: 7074; 1:5000) and anti-mouse (Cat no.: 7076; 1:5000) secondary antibodies were purchased from Cell Signaling Technology. Primary antibodies against CD31 (Abcam, cat no.: ab28364; 1:100), D_2_R (Santa Cruz Biotechnology, cat no.: sc-5303; 1:50), and Ki-67 (Cell Signaling Technology, cat no.: 12202; 1:200) were used for immunofluorescence staining. Fluorescence-conjugated secondary antibody staining was performed in the dark using corresponding Alexa Fluor 488–conjugated anti-rabbit antibody (Molecular Probes; Cat no.: A11008; 1:400) and Alexa Fluor 594–conjugated anti-mouse antibody (Molecular Probes; Cat no.: A11005; 1:400). A mouse monoclonal D_2_R antibody (Santa Cruz Biotechnology, cat no.: sc-5303, 1:50) was used for immunohistochemistry. Cisplatin (232120), etoposide (E1383), quinpirole (Q102), and cabergoline (C0246) were purchased from Sigma-Aldrich.

### Immunoblotting

Stable HUVECs were lysed in SDS containing 2% 1× Laemmli buffer (Bio-Rad) supplemented with 5% 2-mercaptoethanol (Sigma). Cells were immediately scraped off the cell culture plate and transferred to microcentrifuge tubes for boiling. Cell lysates were heated to 95-100 °C for 5 minutes and allowed to cool at room temperature for 10 minutes. Later, 20 µL of cell lysates were separated via 4-20%–gradient SDS-PAGE (Bio-Rad) and transferred to polyvinyl difluoride (PVDF) membranes (Millipore). Following the completion of the protein transfer process, membranes were blocked with 5% bovine serum albumin (Sigma-Aldrich) diluted in 1× Tris-buffered saline (Growcells) containing 0.1% Tween-20 detergent (Fisher Scientific). Membranes were then incubated with primary antibodies overnight at 4°C and corresponding secondary antibodies for 2 h at room temperature. Antibody-reactive protein bands on the membranes were detected in the dark using HRP-reactive chemiluminescence substrate (Thermo Fisher Scientific).

### Immunofluorescence

The tumours of mice bearing chemonaïve and chemoresistant human SCLC PDXs were harvested and fixed in neutral buffered 10% formalin (Sigma) at room temperature for 24 h before processing, embedding in paraffin, and sectioning. Tissues sections were deparaffinized, rehydrated, heat-retrieved with 1× Rodent Decloaker buffer (Biocare Medical, RD913L), and then incubated in Rodent Block M (Biocare Medical, RBM961G) for 1 h at room temperature to eliminate non-specific mouse IgG staining. Tissue sections were then incubated with primary antibodies overnight at 4°C followed by fluorophore-conjugated secondary antibodies for 1 h at room temperature. For the TUNEL staining experiments, tissue sections were rinsed with 1× PBS (Sigma) following incubation with secondary antibodies and subjected to fluorescence-based TUNEL staining following the manufacturer’s protocols (Promega, G3250). Briefly, the tissue specimens were incubated with 1× equilibration buffer for 5 minutes and then with the reaction buffer containing recombinant terminal deoxynucleotidyl transferase (rTdT) enzyme and nucleotide mix for 1 h at 37 °C. The reaction was terminated with 2× saline sodium citrate buffer (SSC). Cell nuclei were counterstained with 4’, 6-diamidino-2-phenylindole, dihydrochloride (DAPI) in ProLong^®^ Gold Antifade Reagent (Cell Signaling Technology). Images were captured using a Zeiss Apotome.2 microscope (20× objective, 0.75 NA) and processed with ZEN microscope software (Zeiss). Double-positive cells (i.e., either TUNEL^+^-CD31^+^ or D_2_R^+^-CD31^+^) per visual field were counted with the cell counter tool available in ImageJ software. The bar graphs were generated by dividing the number of double-positive cells by the number of CD31^+^ blood vessels. The percentage of Ki-67^+^ cells was determined by following the equation: percentage of Ki-67^+^ cells = (Count of Ki-67^+^ cells / Count of DAPI^+^ cells) × 100.

### Immunohistochemistry

We obtained chemonaïve (i.e., before the start of chemotherapy) and matched chemoresistant (i.e., recurrence of tumour after chemotherapy) human SCLC whole-tissue specimens from nine SCLC patients at Mayo Clinic in Rochester, MN, in accordance with institutional review board–approved protocols. Formalin-fixed, paraffin-embedded whole tissues were serially sectioned, mounted on glass slides, and immunostained using primary and HRP-conjugated secondary antibodies. The staining was performed by using a Bond Autostainer (Leica), and the sections were incubated in hematoxylin (IHC World) to detect nuclei. A pulmonary pathologist (Y-C.L.) scored each lung tumour specimen of each group by observing the prevalence of D_2_R staining in the tumour-associated endothelial cells under the light microscope.

### *In vivo* orthotopic lung cancer model

Eight- to ten-week-old pathogen-free SCID/NCr mice (Strain Code: 561) purchased from Charles River Laboratories were bred and maintained in accordance with protocols approved by the University of Minnesota Institutional Animal Care and Use Committee (IACUC). One million luciferase-labelled human DMS-53 SCLC cells suspended in 80 µL PBS (Corning) and high-concentration Matrigel (Corning; Cat. no.: 354248) were orthotopically injected into the left thoracic cavity of 8- to 12-week-old male and female mice anesthetized with pharmaceutical-grade ketamine (90–120 mg/kg) and xylazine (5–10 mg/kg). Bioluminescence imaging of mice anesthetized with isoflurane was performed on the indicated days using the IVIS^®^ Lumina^™^ S5 high-throughput 2D optical imaging system (Perkin Elmer) to monitor lung tumour growth non-invasively.

### Human SCLC PDX subcutaneous mouse model

Ten- to twelve-week-old male and female NOD.Cg-*Prkdc^scid^ Il2rg^tm1Wjl^*/SzJ mice (NSG, Jackson Laboratory, Strain Code: 5557) were bred, maintained in a temperature-controlled room with alternating 12-hour light/dark cycles under specific pathogen–free conditions, and fed a standard diet in accordance with protocols approved by the University of Minnesota IACUC. Male and female mice anesthetized with pharmaceutical-grade ketamine (90–120 mg/kg) and xylazine (5–10 mg/kg) via intraperitoneal injection under laminar flow hood in a specific pathogen–free room were subcutaneously injected with 5 × 10^6^ human SCLC cells collected from either chemonaïve or chemoresistant human SCLC PDX tissue samples. Briefly, frozen vials of tissues from one chemonaïve and three chemoresistant SCLC PDX–bearing mice were transported to our laboratory overnight on dry ice from our collaborator at Memorial Sloan Kettering Cancer Center. Due to the small amount of material available, the entire tumour sample was first resuspended in 100 µL high-concentration Matrigel on ice and later injected subcutaneously in the flanks of anesthetized NSG mice. Tumour growth was monitored weekly. When tumour volume reached 200 mm^3^ in size, mice were randomly divided into treatment groups such that each group had similar mean tumour volumes. Tumour volume was measured every week using the formula (length × width^2^)/2. When the P_0_ tumours reached 2000 mm^3^ in volume, the mouse was sacrificed, tumour tissue samples were finely minced with sterile razor blades under aseptic conditions, vigorously triturated in Accutase™ cell detachment solution (BD Bioscience, 561527), passed through a 70-µm filter (Corning), centrifuged at 433 × g for 5 mins, and either cryopreserved in 85% RPMI-1640 (Corning)/10% FBS (Millipore)/5% DMSO (MP Biomedicals) for future use or suspended in 100 μL PBS and high-concentration Matrigel on ice (5 × 10^6^ cells per mouse). Cells were then subcutaneously injected into the right flank of NSG mice and were monitored for tumour growth. After establishment of palpable tumours (≥100 mm^3^), mice were randomly divided into groups and administered drugs at the indicated doses. At the endpoint, mice were euthanized by CO_2_ asphyxiation followed by cervical dislocation. Extirpated tumours were photographed, weighed, and preserved in neutral buffered 10% formalin (Sigma) for immunofluorescence analysis.

### Clinical workflow and patient selection

Patients who met the following criteria were enrolled in this study: (1) pathologically confirmed advanced SCLC; (2) defined subtypes based on the expression of transcription factors ASCL1, NEUROD1, and POU2F3; (3) treatment with platinum-based chemotherapy in the first-line setting; and (4) available biopsied tumour samples before and after the chemotherapy. A 20% increase of tumour burden after the completion of chemotherapy was considered as disease progression in accordance with RECIST 1.1. Pathological diagnosis and staging were carried out according to the staging system of the 2021 International Association for the Study of Lung Cancer (9^th^ edition). Written informed consent was obtained from all of the patients prior to inclusion in this study. The Mayo Clinic Institutional Review Board Committee approved this study.

### Statistics

To compare differences between two groups, two-way unpaired t-tests were performed and values of *P* ≤ 0.05 were considered significant. A two-way analysis of variance (ANOVA) followed by Sidak’s test was used to determine statistically significant differences between multiple groups (greater than two). Data expressed as mean ± SEM are representative of at least three independent experiments. For most animal experiments, the number of animals per group was calculated based on a one-way ANOVA analysis to allow 90% power when the mean in the test group is 1.25 standard deviations higher or lower than the mean in the controls.

## Acknowledgements

This work was supported by a Research Scholar Grant RSG-21-034-01-TBG from the American Cancer Society and The Hormel Foundation to L.H.H. The generation of the patient-derived xenograft models of chemotherapy-refractory small cell lung cancer was supported by R35 CA263816 and U24 CA213274 to C.M.R. We thank The Hormel Institute and its staff for administrative, shared equipment, animal facility, and institutional support. We are grateful to Dr. Naomi Ruff for providing editorial support.

## Author Contributions

S.K.A. conducted most of the in vitro cell line-based experiments, including immunofluorescence studies, generation of stable cell lines, immunoblotting, etc. A.P. performed the organoid culture and cleaved caspase 3 assays. S.K.A., A.P., L.W., and L.H.H. managed the mouse colony and performed tumour studies in mice. S.K.A., A.P., and L.W. conducted murine in vivo imaging, tumour measurements, and necropsy. P.M. and C.M.R. generated and provided the patient-derived xenograft models of chemotherapy-refractory small cell lung cancer. M.C.A. and Y-C.L. coordinated the acquisition of small cell lung cancer specimens from patients at Mayo Clinic in accordance with IRB-approved protocols. B.A.T. captured images of the small cell lung cancer specimens that had been immunostained to detect D_2_R protein. Y-C.L. led the immunostaining, imaging, pathological review, and analysis of patient-derived lung tumour specimens. S.K.A., A.P., L.W., C.M.R., Y-C.L., and L.H.H. provided technical and scientific support. S.K.A., A.P., and L.H.H. performed experimental troubleshooting, reviewed relevant scientific literature, critically analyzed data, prepared figures, and wrote the manuscript. L.H.H. conceived the aims, led the project, and acquired funding to complete the reported research.

---

**Supplementary Figure 1:**
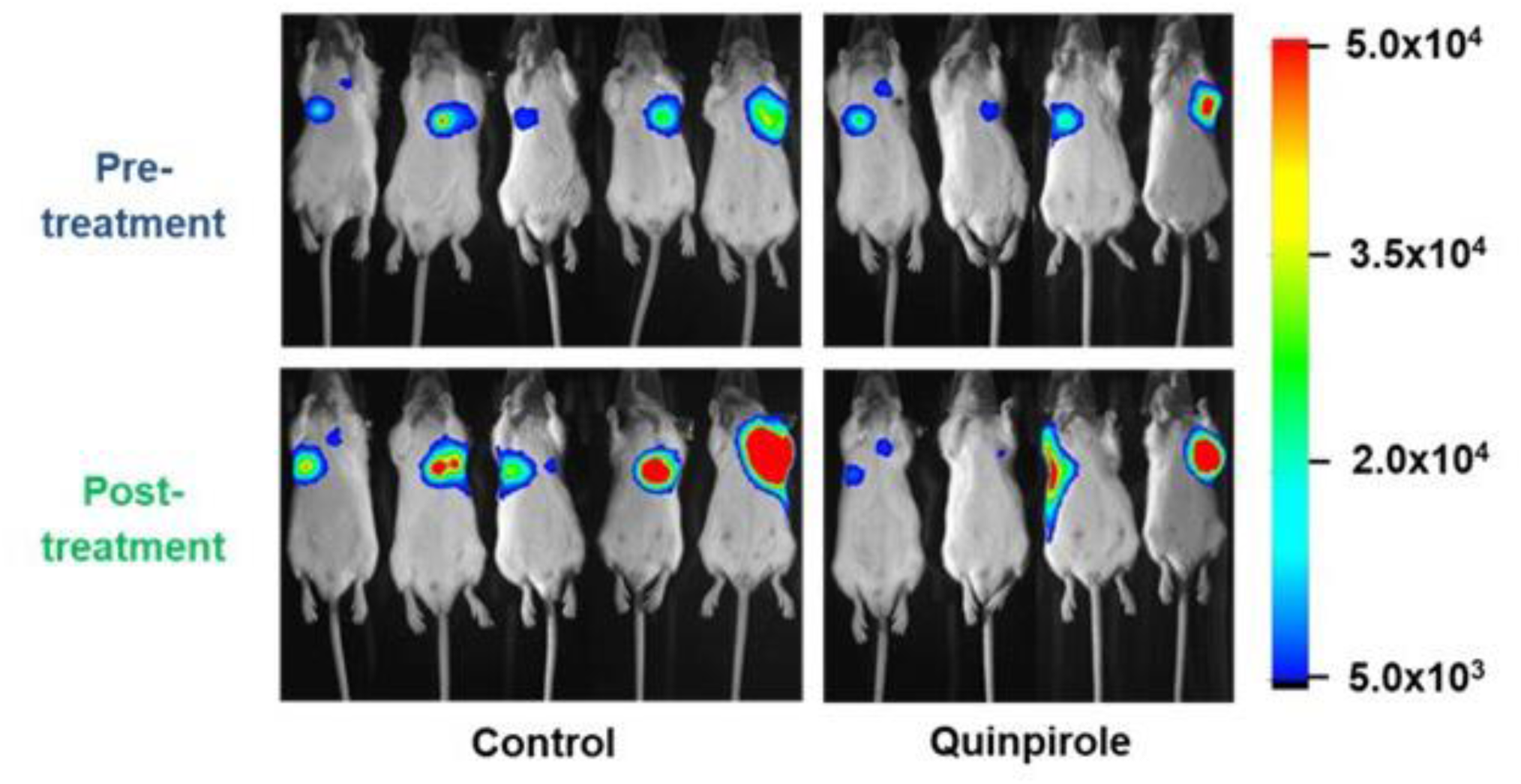
Pre- and post-treatment luminescence images of vehicle- and quinpirole-treated SCID mice. One million luciferase-labelled human DMS-53 SCLC cells were orthotopically injected into the left thoracic cavity of SCID mice. Mice administered either vehicle control (1× PBS) or quinpirole (10 mg/kg) were imaged for bioluminescence before and after the treatment.

**Supplementary Figure 2:**
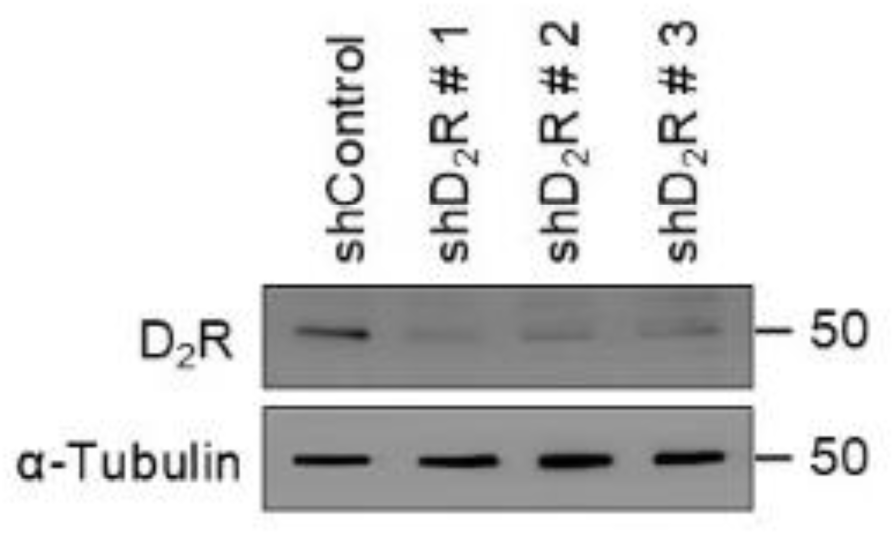
Validation of D_2_R knockdown in endothelial cells. HUVECs transduced with lentivirus encoding either a D_2_R shRNA (#1-3) or control shRNA were lysed. Equal amounts of protein were separated in a 4-20% SDS-PAGE gel followed by protein transfer to PVDF membrane. Antibody-reactive bands were detected using primary antibodies against D_2_R and α-tubulin (loading control).

**Supplementary Figure 3:**
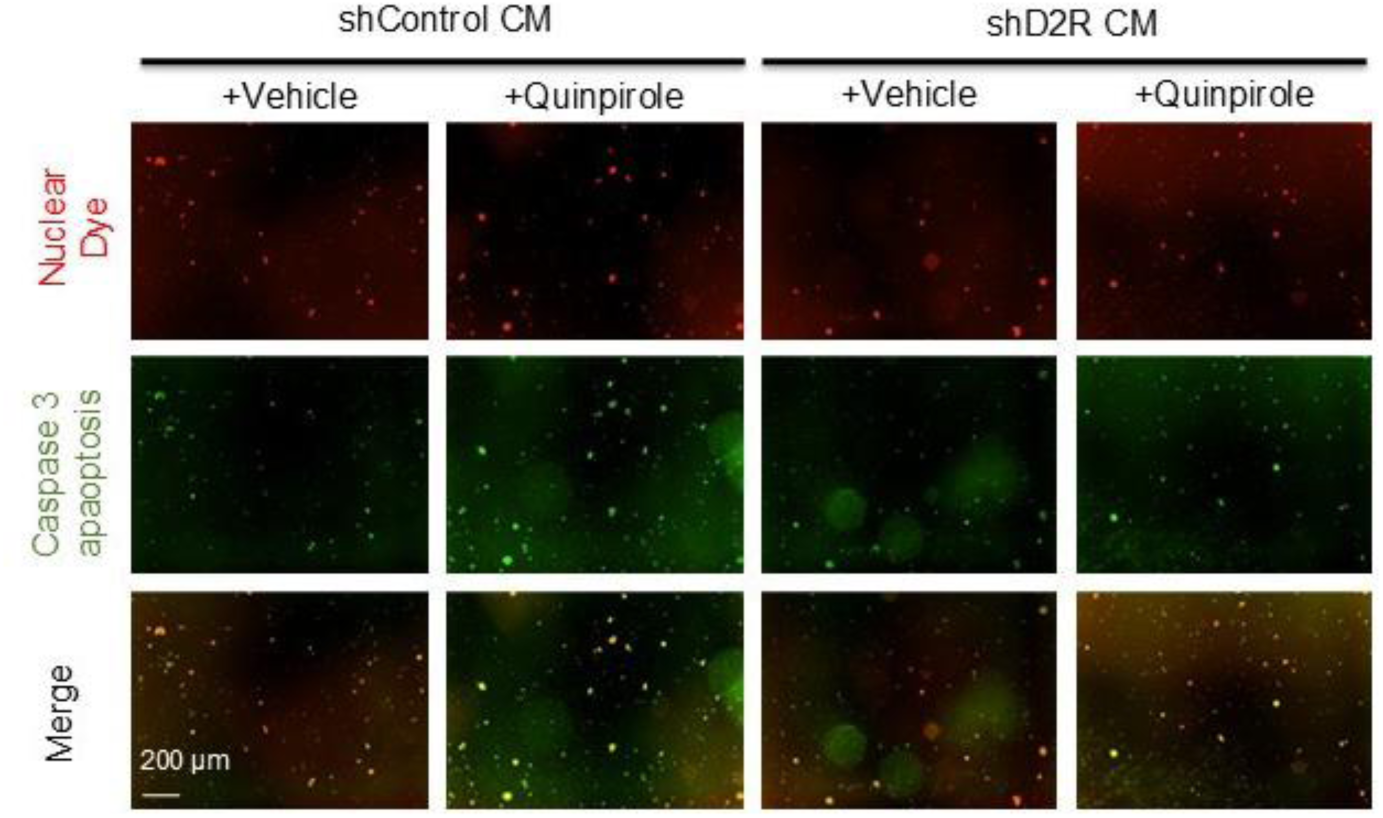
Conditioned medium from human endothelial cells treated with a D_2_R agonist increases caspase-3–mediated apoptosis of SCLC chemotherapy-resistant organoids. SCLC PDX cultured as three-dimensional organoids showed increased apoptosis upon treatment with conditioned media collected from quinpirole (50 µM)-treated HUVECs.

**Supplementary Figure 4:**
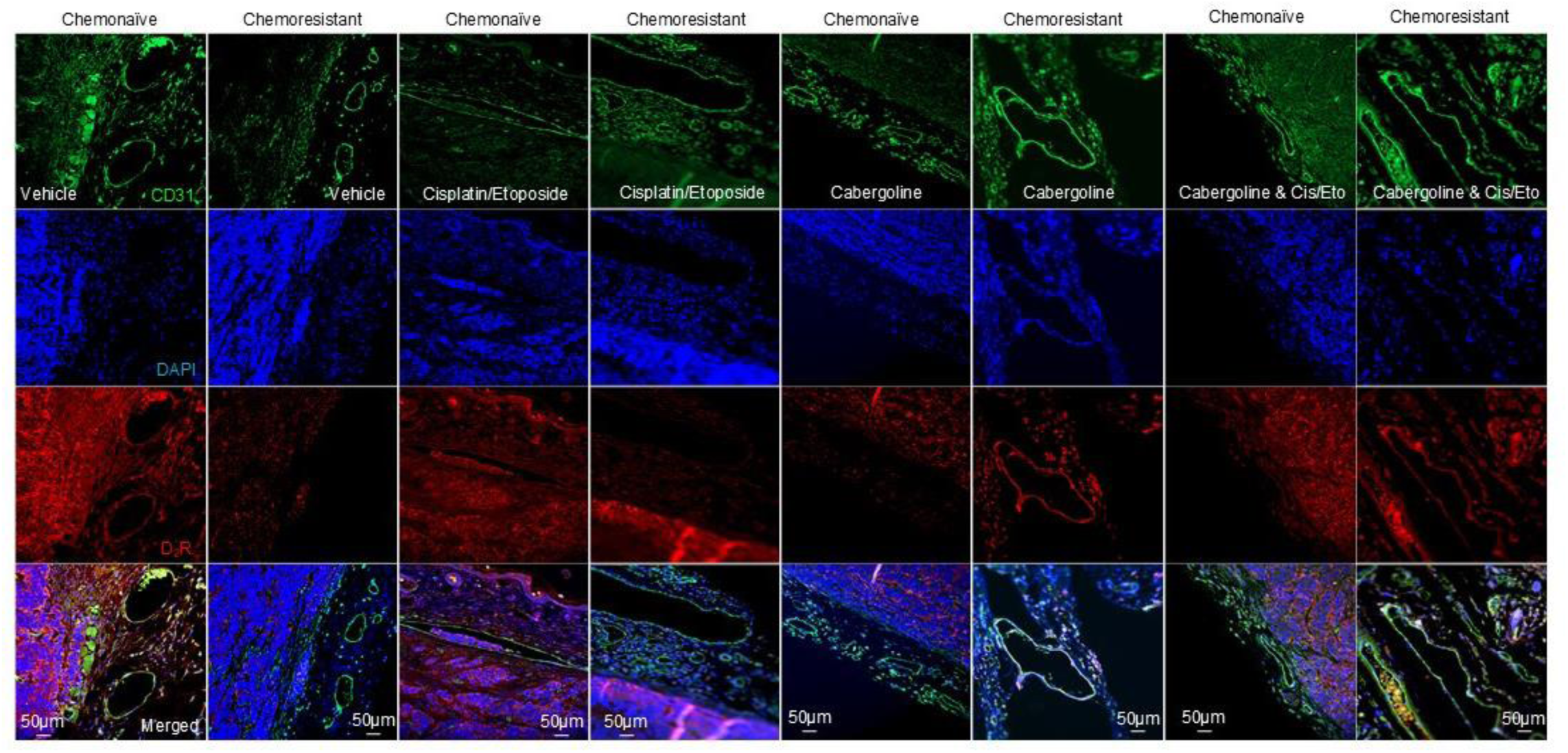
Co-immunofluorescence images of D_2_R and CD31 immunostaining in chemonaïve and chemoresistant SCLC PDXs. NSG mice were subcutaneously implanted with 5 × 10^6^ cells obtained from either chemonaïve (MSK-LX40) or chemoresistant (MSK-LX40R) human SCLC PDXs. Mice were randomly divided into four groups to receive 1) vehicle; 2) cisplatin/etoposide; 3) cabergoline; or 4) cabergoline and cisplatin/etoposide. At the endpoint, mice were sacrificed, tumours were resected, and co-immunofluorescence staining was performed on FFPE tumour tissues (n=3 per group) using primary antibodies against CD31 and D_2_R. Nuclei were counterstained with DAPI.

**Supplementary Table 1:** D_2_R protein expression in tumour-associated endothelial cells in paired chemotherapy-naïve and chemotherapy-resistant patient specimens.

| **Patient Number** | **Subtype** | **Chemotherapy Status** | **D<sub>2</sub>R Staining in Endothelium** |
| --- | --- | --- | --- |
| 4 | SCLC-A | Chemotherapy-naïve | 40% |
| 4 | SCLC-A | Chemotherapy-resistant | 1% |
| 7 | SCLC-A | Chemotherapy-naïve | 1% |
| 7 | SCLC-A | Chemotherapy-resistant | 30% |
| 16 | SCLC-A | Chemotherapy-naïve | 20% |
| 16 | SCLC-A | Chemotherapy-resistant | 5% |
| 17 | SCLC-A | Chemotherapy-naïve | 50% |
| 17 | SCLC-A | Chemotherapy-resistant | 30% |
| 22 | SCLC-A | Chemotherapy-naïve | 40% |
| 22 | SCLC-A | Chemotherapy-resistant | 60% |
| 25 | SCLC-A | Chemotherapy-naïve | 30% |
| 25 | SCLC-A | Chemotherapy-resistant | 1% |
| 29 | SCLC-A | Chemotherapy-naïve | 50% |
| 29 | SCLC-A | Chemotherapy-resistant | 5% |
| 32 | SCLC-A | Chemotherapy-naïve | 20% |
| 32 | SCLC-A | Chemotherapy-resistant | 10% |
| 34 | SCLC-A | Chemotherapy-naïve | 50% |
| 34 | SCLC-A | Chemotherapy-resistant | 1% |

## References

1 Bray, F. et al. Global cancer statistics 2018: GLOBOCAN estimates of incidence and mortality worldwide for 36 cancers in 185 countries. CA: a cancer journal for clinicians (2018). 10.3322/caac.21492

2 Kahnert, K., Kauffmann-Guerrero, D. & Huber, R. M. SCLC-State of the Art and What Does the Future Have in Store? Clinical lung cancer (2016). 10.1016/j.cllc.2016.05.014

3 Riaz, S. P. et al. Trends in incidence of small cell lung cancer and all lung cancer. Lung Cancer 75, 280–284 (2012). 10.1016/j.lungcan.2011.08.004

4 Kalemkerian, G. P. et al. Small cell lung cancer. Journal of the National Comprehensive Cancer Network: JNCCN 11, 78–98 (2013).

5 Miller, K. D. et al. Cancer treatment and survivorship statistics, 2016. CA: a cancer journal for clinicians 66, 271–289 (2016). 10.3322/caac.21349

6 Gay, C. M. et al. Patterns of transcription factor programs and immune pathway activation define four major subtypes of SCLC with distinct therapeutic vulnerabilities. Cancer Cell 39, 346–360 e347 (2021). 10.1016/j.ccell.2020.12.014

7 Rudin, C. M. et al. Molecular subtypes of small cell lung cancer: a synthesis of human and mouse model data. Nat Rev Cancer 19, 289–297 (2019). 10.1038/s41568-019-0133-9

8 McNamee, N., da Silva, I. P., Nagrial, A. & Gao, B. Small-Cell Lung Cancer-An Update on Targeted and Immunotherapies. Int J Mol Sci 24 (2023). 10.3390/ijms24098129

9 Rossi, A. et al. Carboplatin- or cisplatin-based chemotherapy in first-line treatment of small-cell lung cancer: the COCIS meta-analysis of individual patient data. Journal of clinical oncology : official journal of the American Society of Clinical Oncology 30, 1692–1698 (2012). 10.1200/JCO.2011.40.4905

10 Goldman, J. W. et al. Durvalumab, with or without tremelimumab, plus platinum-etoposide versus platinum-etoposide alone in first-line treatment of extensive-stage small-cell lung cancer (CASPIAN): updated results from a randomised, controlled, open-label, phase 3 trial. The Lancet. Oncology 22, 51–65 (2021). 10.1016/S1470-2045(20)30539-8

11 Liu, S. V. et al. Updated Overall Survival and PD-L1 Subgroup Analysis of Patients With Extensive-Stage Small-Cell Lung Cancer Treated With Atezolizumab, Carboplatin, and Etoposide (IMpower133). Journal of clinical oncology : official journal of the American Society of Clinical Oncology 39, 619–630 (2021). 10.1200/JCO.20.01055

12 Senger, D. R. et al. Tumor cells secrete a vascular permeability factor that promotes accumulation of ascites fluid. Science 219, 983–985 (1983).

13 Leung, D. W., Cachianes, G., Kuang, W. J., Goeddel, D. V. & Ferrara, N. Vascular endothelial growth factor is a secreted angiogenic mitogen. Science 246, 1306–1309 (1989).

14 Carmeliet, P. et al. Abnormal blood vessel development and lethality in embryos lacking a single VEGF allele. Nature 380, 435–439 (1996). 10.1038/380435a0

15 Ferrara, N. et al. Heterozygous embryonic lethality induced by targeted inactivation of the VEGF gene. Nature 380, 439–442 (1996). 10.1038/380439a0

16 Nasevicius, A., Larson, J. & Ekker, S. C. Distinct requirements for zebrafish angiogenesis revealed by a VEGF-A morphant. Yeast 17, 294–301 (2000). 10.1002/1097-0061(200012)17:4<294::AID-YEA54>3.0.CO;2-5

17 Olsson, A. K., Dimberg, A., Kreuger, J. & Claesson-Welsh, L. VEGF receptor signalling - in control of vascular function. Nat Rev Mol Cell Biol 7, 359–371 (2006). 10.1038/nrm1911

18 Simons, M., Gordon, E. & Claesson-Welsh, L. Mechanisms and regulation of endothelial VEGF receptor signalling. Nat Rev Mol Cell Biol 17, 611–625 (2016). 10.1038/nrm.2016.87

19 Kaner, R. J. et al. Lung overexpression of the vascular endothelial growth factor gene induces pulmonary edema. Am J Respir Cell Mol Biol 22, 657–664 (2000).

20 Lahm, T. et al. The critical role of vascular endothelial growth factor in pulmonary vascular remodeling after lung injury. Shock 28, 4–14 (2007). 10.1097/shk.0b013e31804d1998

21 Lin, Q. et al. Prognostic value of serum IL-17 and VEGF levels in small cell lung cancer. The International journal of biological markers 30, e359–363 (2015). 10.5301/jbm.5000148

22 Nowak, K. et al. Circulating endothelial progenitor cells are increased in human lung cancer and correlate with stage of disease. European journal of cardio-thoracic surgery : official journal of the European Association for Cardio-thoracic Surgery 37, 758–763 (2010). 10.1016/j.ejcts.2009.10.002

23 Salven, P., Ruotsalainen, T., Mattson, K. & Joensuu, H. High pre-treatment serum level of vascular endothelial growth factor (VEGF) is associated with poor outcome in small-cell lung cancer. International journal of cancer 79, 144–146 (1998).

24 Zhan, P. et al. Prognostic value of vascular endothelial growth factor expression in patients with lung cancer: a systematic review with meta-analysis. Journal of thoracic oncology : official publication of the International Association for the Study of Lung Cancer 4, 1094–1103 (2009). 10.1097/JTO.0b013e3181a97e31

25 Jenab-Wolcott, J. & Giantonio, B. J. Bevacizumab: current indications and future development for management of solid tumors. Expert opinion on biological therapy 9, 507–517 (2009). 10.1517/14712590902817817

26 Limaverde-Sousa, G., Sternberg, C. & Ferreira, C. G. Antiangiogenesis beyond VEGF inhibition: a journey from antiangiogenic single-target to broad-spectrum agents. Cancer treatment reviews 40, 548–557 (2014). 10.1016/j.ctrv.2013.11.009

27 Aggarwal, C., Somaiah, N. & Simon, G. Antiangiogenic agents in the management of non-small cell lung cancer: where do we stand now and where are we headed? Cancer biology & therapy 13, 247–263 (2012). 10.4161/cbt.19594

28 Horn, L. et al. Phase II study of cisplatin plus etoposide and bevacizumab for previously untreated, extensive-stage small-cell lung cancer: Eastern Cooperative Oncology Group Study E3501. Journal of clinical oncology : official journal of the American Society of Clinical Oncology 27, 6006–6011 (2009). 10.1200/JCO.2009.23.7545

29 Mountzios, G. et al. Paclitaxel plus bevacizumab in patients with chemoresistant relapsed small cell lung cancer as salvage treatment: a phase II multicenter study of the Hellenic Oncology Research Group. Lung cancer 77, 146–150 (2012). 10.1016/j.lungcan.2012.02.002

30 Petrioli, R. et al. Cisplatin, Etoposide, and Bevacizumab Regimen Followed by Oral Etoposide and Bevacizumab Maintenance Treatment in Patients With Extensive -Stage Small Cell Lung Cancer: A Single-Institution Experience. Clinical lung cancer 16, e229–234 (2015). 10.1016/j.cllc.2015.05.005

31 Pujol, J. L. et al. Randomized phase II-III study of bevacizumab in combination with chemotherapy in previously untreated extensive small-cell lung cancer: results from the IFCT-0802 trialdagger. Annals of oncology : official journal of the European Society for Medical Oncology / ESMO 26, 908–914 (2015). 10.1093/annonc/mdv065

32 Ready, N. E. et al. Cisplatin, irinotecan, and bevacizumab for untreated extensive-stage small-cell lung cancer: CALGB 30306, a phase II study. Journal of clinical oncology : official journal of the American Society of Clinical Oncology 29, 4436–4441 (2011). 10.1200/JCO.2011.35.6923

33 Spigel, D. R. et al. Phase II trial of irinotecan, carboplatin, and bevacizumab in the treatment of patients with extensive-stage small-cell lung cancer. Journal of thoracic oncology : official publication of the International Association for the Study of Lung Cancer 4, 1555–1560 (2009). 10.1097/JTO.0b013e3181bbc540

34 Spigel, D. R. et al. Randomized phase II study of bevacizumab in combination with chemotherapy in previously untreated extensive-stage small-cell lung cancer: results from the SALUTE trial. Journal of clinical oncology : official journal of the American Society of Clinical Oncology 29, 2215–2222 (2011). 10.1200/JCO.2010.29.3423

35 Spigel, D. R., Waterhouse, D. M., Lane, S., Legenne, P. & Bhatt, K. Efficacy and safety of oral topotecan and bevacizumab combination as second-line treatment for relapsed small-cell lung cancer: an open-label multicenter single-arm phase II study. Clinical lung cancer 14, 356–363 (2013). 10.1016/j.cllc.2012.12.003

36 Lee, S. M. et al. Anti-angiogenic therapy using thalidomide combined with chemotherapy in small cell lung cancer: a randomized, double-blind, placebo-controlled trial. Journal of the National Cancer Institute 101, 1049–1057 (2009). 10.1093/jnci/djp200

37 Pujol, J. L. et al. Phase III double-blind, placebo-controlled study of thalidomide in extensive-disease small-cell lung cancer after response to chemotherapy: an intergroup study FNCLCC cleo04 IFCT 00-01. Journal of clinical oncology : official journal of the American Society of Clinical Oncology 25, 3945–3951 (2007). 10.1200/JCO.2007.11.8109

38 Han, J. Y. et al. A phase II study of sunitinib in patients with relapsed or refractory small cell lung cancer. Lung cancer 79, 137–142 (2013). 10.1016/j.lungcan.2012.09.019

39 Ready, N. E. et al. Chemotherapy With or Without Maintenance Sunitinib for Untreated Extensive -Stage Small-Cell Lung Cancer: A Randomized, Double-Blind, Placebo-Controlled Phase II Study-CALGB 30504 (Alliance). Journal of clinical oncology : official journal of the American Society of Clinical Oncology 33, 1660–1665 (2015). 10.1200/JCO.2014.57.3105

40 Schneider, B. J. et al. Phase II trial of sunitinib maintenance therapy after platinum-based chemotherapy in patients with extensive-stage small cell lung cancer. Journal of thoracic oncology : official publication of the International Association for the Study of Lung Cancer 6, 1117–1120 (2011). 10.1097/JTO.0b013e31821529c3

41 Spigel, D. R. et al. Phase II study of maintenance sunitinib following irinotecan and carboplatin as first-line treatment for patients with extensive-stage small-cell lung cancer. Lung cancer 77, 359–364 (2012). 10.1016/j.lungcan.2012.03.009

42 Arnold, A. M. et al. Phase II study of vandetanib or placebo in small-cell lung cancer patients after complete or partial response to induction chemotherapy with or without radiation therapy: National Cancer Institute of Canada Clinical Trials Group Study BR.20. Journal of clinical oncology : official journal of the American Society of Clinical Oncology 25, 4278–4284 (2007). 10.1200/JCO.2007.12.3083

43 Ramalingam, S. S. et al. Phase II study of Cediranib (AZD 2171), an inhibitor of the vascular endothelial growth factor receptor, for second-line therapy of small cell lung cancer (National Cancer Institute #7097). Journal of thoracic oncology : official publication of the International Association for the Study of Lung Cancer 5, 1279–1284 (2010). 10.1097/JTO.0b013e3181e2fcb0

44 Gitlitz, B. J. et al. Sorafenib in platinum-treated patients with extensive stage small cell lung cancer: a Southwest Oncology Group (SWOG 0435) phase II trial. Journal of thoracic oncology : official publication of the International Association for the Study of Lung Cancer 5, 1835–1840 (2010). 10.1097/JTO.0b013e3181f0bd78

45 Han, J. Y., Kim, H. Y., Lim, K. Y., Hwangbo, B. & Lee, J. S. A phase II study of nintedanib in patients with relapsed small cell lung cancer. Lung cancer 96, 108–112 (2016). 10.1016/j.lungcan.2016.04.002

46 Shi, J. et al. Anlotinib as third- or further-line therapy for short-term relapsed small-cell lung cancer: subgroup analysis of a randomized phase 2 study (ALTER1202). Front Med 16, 766–772 (2022). 10.1007/s11684-021-0916-8

47 Song, P. F., Xu, N. & Li, Q. Efficacy and Safety of Anlotinib for Elderly Patients with Previously Treated Extensive-Stage SCLC and the Prognostic Significance of Common Adverse Reactions. Cancer Manag Res 12, 11133–11143 (2020). 10.2147/CMAR.S275624

48 Zhang, X., Liu, Q., Liao, Q. & Zhao, Y. Potential Roles of Peripheral Dopamine in Tumor Immunity. Journal of Cancer 8, 2966–2973 (2017). 10.7150/jca.20850

49 Chakroborty, D., Sarkar, C., Basu, B., Dasgupta, P. S. & Basu, S. Catecholamines regulate tumor angiogenesis. Cancer research 69, 3727–3730 (2009). 10.1158/0008-5472.CAN-08-4289

50 Bhattacharya, R. et al. The neurotransmitter dopamine modulates vascular permeability in the endothelium. J Mol Signal 3, 14 (2008). 10.1186/1750-2187-3-14

51 Sinha, S. et al. Dopamine regulates phosphorylation of VEGF receptor 2 by engaging Src-homology-2- domain-containing protein tyrosine phosphatase 2. J Cell Sci 122, 3385–3392 (2009). 10.1242/jcs.053124

52 Basu, S. et al. The neurotransmitter dopamine inhibits angiogenesis induced by vascular permeability factor/vascular endothelial growth factor. Nature medicine 7, 569–574 (2001). 10.1038/87895

53 Basu, S. et al. Ablation of peripheral dopaminergic nerves stimulates malignant tumor growth by inducing vascular permeability factor/vascular endothelial growth factor-mediated angiogenesis. Cancer research 64, 5551–5555 (2004). 10.1158/0008-5472.CAN-04-1600

54 Sarkar, C., Chakroborty, D., Chowdhury, U. R., Dasgupta, P. S. & Basu, S. Dopamine increases t he efficacy of anticancer drugs in breast and colon cancer preclinical models. Clinical cancer research : an official journal of the American Association for Cancer Research 14, 2502–2510 (2008). 10.1158/1078-0432.CCR-07-1778

55 Alam, S. K. et al. DARPP-32 and t-DARPP promote non-small cell lung cancer growth through regulation of IKKalpha-dependent cell migration. Communications biology 1, 43 (2018). 10.1038/s42003-018-0050-6

56 Alam, S. K. et al. ASCL1-regulated DARPP-32 and t-DARPP stimulate small cell lung cancer growth and neuroendocrine tumour cell proliferation. British journal of cancer (2020). 10.1038/s41416-020-0923-6

57 Alam, S. K., Wang, L., Zhu, Z. & Hoeppner, L. H. IKKalpha promotes lung adenocarcinoma growth through ERK signaling activation via DARPP-32-mediated inhibition of PP1 activity. NPJ Precis Oncol 7, 33 (2023). 10.1038/s41698-023-00370-3

58 Alam, S. K. et al. DARPP-32 promotes ERBB3-mediated resistance to molecular targeted therapy in EGFR-mutated lung adenocarcinoma. Oncogene 41, 83–98 (2022). 10.1038/s41388-021-02028-5

59 Vohra, P. K. et al. Dopamine inhibits pulmonary edema through the VEGF-VEGFR2 axis in a murine model of acute lung injury. Am J Physiol Lung Cell Mol Physiol 302, L185–192 (2012). 10.1152/ajplung.00274.2010

60 Hoeppner, L. H. et al. Dopamine D2 receptor agonists inhibit lung cancer progression by reducing angiogenesis and tumor infiltrating myeloid derived suppressor cells. Mol Oncol 9, 270–281 (2015). 10.1016/j.molonc.2014.08.008

61 Chakroborty, D. et al. Dopamine regulates endothelial progenitor cell mobilization from mouse bone marrow in tumor vascularization. J Clin Invest 118, 1380–1389 (2008). 10.1172/JCI33125

62 Tilan, J. & Kitlinska, J. Sympathetic Neurotransmitters and Tumor Angiogenesis -Link between Stress and Cancer Progression. J Oncol 2010, 539706 (2010). 10.1155/2010/539706

63 Gardner, E. E. et al. Chemosensitive Relapse in Small Cell Lung Cancer Proceeds through an EZH2- SLFN11 Axis. Cancer Cell 31, 286–299 (2017). 10.1016/j.ccell.2017.01.006

64 Jain, R. K. Normalization of tumor vasculature: an emerging concept in antiangiogenic therapy. Science 307, 58–62 (2005). 10.1126/science.1104819

65 Ferrara, N., Hillan, K. J., Gerber, H. P. & Novotny, W. Discovery and development of bevacizumab, an anti-VEGF antibody for treating cancer. Nature reviews. Drug discovery 3, 391–400 (2004). 10.1038/nrd1381

66 Ye, W. The Complexity of Translating Anti-angiogenesis Therapy from Basic Science to the Clinic. Developmental cell 37, 114–125 (2016). 10.1016/j.devcel.2016.03.015

67 Cohen, M. H., Shen, Y. L., Keegan, P. & Pazdur, R. FDA drug approval summary: bevacizumab (Avastin) as treatment of recurrent glioblastoma multiforme. Oncologist 14, 1131–1138 (2009). 10.1634/theoncologist.2009-0121

68 Pujol, J. L. et al. Randomized phase II–III study of bevacizumab in combination with chemotherapy in previously untreated extensive small-cell lung cancer: results from the IFCT-0802 trial†. Annals of Oncology 26, 908–914 (2015). 10.1093/annonc/mdv065

69 Jalal, S. et al. Paclitaxel Plus Bevacizumab in Patients with Chemosensitive Relapsed Small Cell Lung Cancer: A Safety, Feasibility, and Efficacy Study from the Hoosier Oncology Group. Journal of Thoracic Oncology 5, 2008–2011 (2010). 10.1097/JTO.0b013e3181f77b6e

70 Sandler, A. et al. Paclitaxel-carboplatin alone or with bevacizumab for non-small-cell lung cancer. The New England journal of medicine 355, 2542–2550 (2006). 10.1056/NEJMoa061884

71 Grant, C. E., Flis, A. L. & Ryan, B. M. Understanding the role of dopamine in cancer: past, present and future. Carcinogenesis 43, 517–527 (2022). 10.1093/carcin/bgac045

72 Ito, T. et al. Pulmonary Neuroendocrine Cells and Small Cell Lung Carcinoma: Immunohistochemical Study Focusing on Mechanisms of Neuroendocrine Differentiation. Acta Histochem Cytochem 55, 75–83 (2022). 10.1267/ahc.22-00031

73 Xie, X., Luo, X., Xie, M., Liu, Y. & Wu, T. Risk of lung cancer in Parkinson’s disease. Oncotarget 7, 77319–77325 (2016). 10.18632/oncotarget.12964

74 Melmed, S. et al. Diagnosis and treatment of hyperprolactinemia: an Endocrine Society clinical practice guideline. J Clin Endocrinol Metab 96, 273–288 (2011). 10.1210/jc.2010-1692

75 Auriemma, R. S., Pirchio, R., De Alcubierre, D., Pivonello, R. & Colao, A. Dopamine Agonists: From the 1970s to Today. Neuroendocrinology 109, 34–41 (2019). 10.1159/000499470

76 Sarkar, C. et al. Dopamine in vivo inhibits VEGF-induced phosphorylation of VEGFR-2, MAPK, and focal adhesion kinase in endothelial cells. Am J Physiol Heart Circ Physiol 287, H1554–1560 (2004). 10.1152/ajpheart.00272.2004

77 Moreno-Smith, M. et al. Dopamine blocks stress-mediated ovarian carcinoma growth. Clin Cancer Res 17, 3649–3659 (2011). 10.1158/1078-0432.Ccr-10-2441

78 Chakroborty, D. et al. Dopamine stabilizes tumor blood vessels by up-regulating angiopoietin 1 expression in pericytes and Kruppel-like factor-2 expression in tumor endothelial cells. Proc Natl Acad Sci U S A 108, 20730–20735 (2011). 10.1073/pnas.1108696108

79 Huang, Y. et al. Vascular normalizing doses of antiangiogenic treatment reprogram the immunosuppressive tumor microenvironment and enhance immunotherapy. Proc Natl Acad Sci U S A 109, 17561–17566 (2012). 10.1073/pnas.1215397109

80 Allen, E. et al. Combined antiangiogenic and anti-PD-L1 therapy stimulates tumor immunity through HEV formation. Sci Transl Med 9 (2017). 10.1126/scitranslmed.aak9679

